# Antipsychotic drugs selectively decorrelate long-range interactions in deep cortical layers

**DOI:** 10.1101/2022.01.31.478462

**Authors:** Matthias Heindorf, Georg B. Keller

## Abstract

Psychosis is characterized by a diminished ability of the brain to distinguish externally driven activity patterns from self-generated activity patterns. Antipsychotic drugs are a class of small molecules with relatively broad binding affinity for a variety of neuromodulator receptors that, in humans, can prevent or ameliorate psychosis. How these drugs influence the function of cortical circuits, and in particular their ability to distinguish between externally and self-generated activity patterns, is still largely unclear. To have experimental control over self-generated sensory feedback we used a virtual reality environment in which the coupling between movement and visual feedback can be altered. We then used widefield calcium imaging to determine the cell type specific functional effects of antipsychotic drugs in mouse dorsal cortex under different conditions of visuomotor coupling. By comparing cell type specific activation patterns between locomotion onsets that were experimentally coupled to self-generated visual feedback and locomotion onsets that were not coupled, we show that deep cortical layers were differentially activated in these two conditions. We then show that the antipsychotic drug clozapine disrupted visuomotor integration at locomotion onsets also primarily in deep cortical layers. Given that one of the key components of visuomotor integration in cortex is long-range cortico-cortical connections, we tested whether the effect of clozapine was detectable in the correlation structure of activity patterns across dorsal cortex. We found that clozapine as well as two other antipsychotic drugs, aripiprazole and haloperidol, resulted in a strong reduction in correlations of layer 5 activity between cortical areas and impaired the spread of visuomotor prediction errors generated in visual cortex. Our results are consistent with the interpretation that a major functional effect of antipsychotic drugs is a selective alteration of long-range layer 5 mediated communication.

## INTRODUCTION

Under normal conditions, self-generated movements are coupled to sensory feedback in different modalities. Locomotion, for example, is coupled to proprioceptive feedback from muscle movements, somatosensory and auditory feedback from touching the ground, vestibular feedback from translational acceleration, and visual feedback in the form of expanding visual flow. This coupling is key to our brain’s ability to learn internal models of the world (Jordan and Rumelhart, 1992). Internal models are the transformations between cortical areas and can be used to predict the sensory consequences of movement. Hallucinations and delusions can be explained as a misconfiguration of these internal models: inaccurate predictions of the sensory consequences of movements can result in self-generated activity that is not correctly distinguished from externally driven activity (Fletcher and Frith, 2009; Frith, 2005).

Self-generated movements result in broad activation of most regions of the brain (Musall et al., 2019; Stringer et al., 2019), including some primary sensory areas like visual cortex (Keller et al., 2012; Saleem et al., 2013) where they have been shown to depend on visual context (Pakan et al., 2016). The framework of predictive processing postulates that in cortex, these movement related signals are internal model based predictions of the sensory consequences of movement that are compared to externally generated bottom-up input to compute prediction errors (Jiang and Rao, 2021; Keller and Mrsic-Flogel, 2018). Evidence for this interpretation has mainly come from the discovery of movement related prediction error responses in a variety of different cortical regions and species (Attinger et al., 2017; Audette et al., 2021; Ayaz et al., 2019; Eliades and Wang, 2008; Heindorf et al., 2018; Keller et al., 2012; Keller and Hahnloser, 2009; Stanley and Miall, 2007; Zmarz and Keller, 2016), and the observation that top-down inputs from motor areas of cortex appear to convey movement related predictions of sensory input to sensory areas of cortex (Leinweber et al., 2017; Schneider et al., 2018). The neurons on which the opposing bottom-up input and top-down prediction converge are referred to as prediction error neurons and are thought to reside primarily in layer 2/3 (L2/3) of cortex (Jordan and Keller, 2020). To complete the circuit, there also needs to be a population of neurons that integrate over prediction error responses. These are referred to as internal representation neurons. It is still unclear which cortical neuron type functions as internal representation neurons, but based on anatomical arguments it has been speculated that they can be found in deep cortical layers (Bastos et al., 2012).

In humans, externally driven and self-generated activity patterns can be experimentally separated at least partially, for example, by showing subjects a visual stimulus and later asking them to visualize the same stimulus (Le Bihan et al., 1993). This type of experiment is considerably more difficult to perform in animals. Using movement as a proxy for self-generated activity patterns, however, using a virtual environment, we can compare a condition in which self-generated and externally generated activity patterns are correlated to a condition in which they are not. In the visual system, we can use a virtual reality environment to experimentally couple or decouple locomotion from visual flow feedback. We will refer to the former as a closed loop condition and to the latter as an open loop condition. Assuming the predictive processing model is correct, we should find differences in the activation patterns in dorsal cortex in response to locomotion onsets as a function of whether they occurred with or without coupled visual flow feedback. Both prediction error neurons as well as internal representation neurons should respond differentially. Thus, we set out to investigate whether there are differences between closed and open loop activation patterns of mouse dorsal cortex, and whether these differences are neuron type specific. If hallucinations are indeed the result of aberrant activation of internal representations, possibly as a result of misconfigured internal models, antipsychotic drugs should alter the differential activation patterns in these neurons.

## RESULTS

### Tlx3 positive layer 5 (L5) intratelencephalic (IT) neurons distinguished closed and open loop visuomotor coupling

To investigate neuron type specific differences in activity patterns at locomotion onsets with and without self-generated visual feedback in mouse dorsal cortex, we quantified locomotion onset activity using widefield calcium imaging (Wekselblatt et al., 2016). A GCaMP variant was expressed using either a retro-orbital injection of an AAV-PHP.eB vector (Chan et al., 2017), or a cross between a Cre line and the Ai148 GCaMP6 reporter line (Daigle et al., 2018) (see Methods; **Figure 1A**, **Table S2**). Mice expressed a GCaMP variant either brain wide (C57BL/6), or in genetically identified subsets of excitatory or inhibitory neurons using Cre driver lines. Please note, we use the term ‘brain wide’ here to describe an expression pattern that was determined by the intersection of the AAV-PHP.eB tropism and the expression pattern of EF1α or hSyn1 promoters but that was otherwise brain wide (see Discussion). Prior to the start of the imaging experiments, mice were implanted with a crystal skull cranial window (Kim et al., 2016). We used crystal skull preparations as we found that hemodynamic artifacts were smaller in this preparation compared to those observed in clear skull preparations (**Figures S1A and S1B**). For all experiments, mice were head fixed and free to locomote on a spherical treadmill (**Figure 1B**). Activity from dorsal cortex was imaged while mice were in a closed loop visuomotor condition, in which locomotion on the spherical treadmill controlled forward movement in a virtual corridor. Following this, mice were exposed to an open loop condition during which mice were presented with a replay of the visual flow they had self-generated in the preceding closed loop condition.

**Figure 1.**
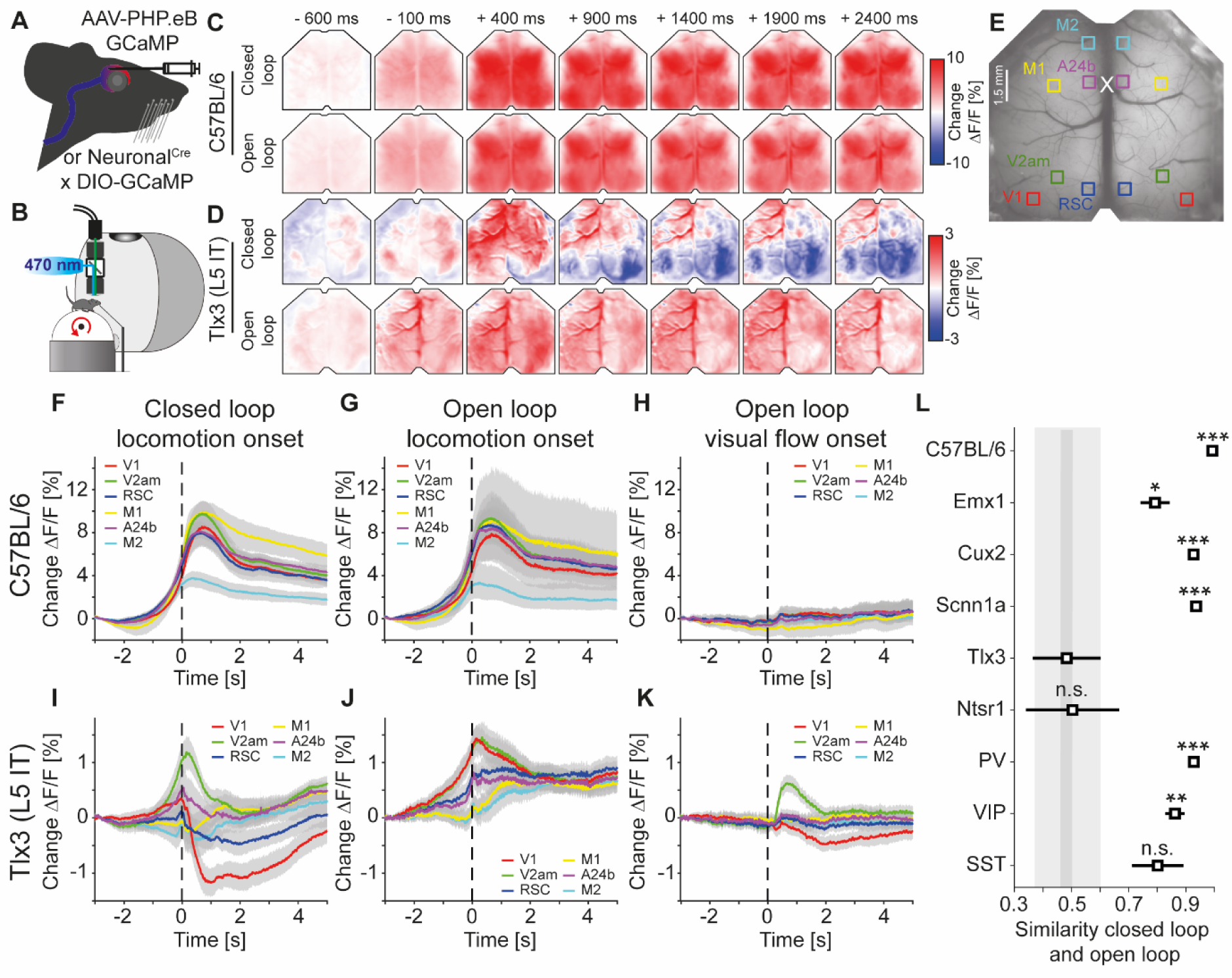
Activation patterns in deep cortical layers distinguished closed and open loop locomotion onsets more strongly than superficial layers. (A) Schematic of GCaMP expression strategy. We either injected an AAV-PHP.eB virus retro-orbitally to express GCaMP brain wide (C57BL/6), in cortical excitatory neurons (Emx1-Cre) or in a subset of SST positive interneurons (see Methods and **Table S2**), or used the progeny of a cross of a cell type specific Cre driver line (Neuronal^Cre^: Cux2-CreERT2, Scnn1a-Cre, Tlx3-Cre, Ntsr1-Cre, PV-Cre, VIP-Cre or SST-Cre) with the Ai148 GCaMP6 reporter line. All mice were then implanted with a crystal skull cranial window prior to imaging experiments. (B) Schematic of the experimental setup. We imaged GCaMP fluorescence under 470 nm LED illumination with an sCMOS camera through a macroscope. Mice were free to locomote on an air supported spherical treadmill while coupled (closed loop), uncoupled (open loop), or no (dark) visual flow in the form of movement along a virtual corridor was displayed on a toroidal screen placed in front of the mouse. Walls of the virtual corridor were patterned with vertical sinusoidal gratings. In a separate condition, we then presented drifting grating stimuli (grating session, see Methods). (C) Average response in an example C57BL/6 mouse that expressed GCaMP6 brain wide during closed loop locomotion onsets (top row, 83 onsets) and open loop locomotion onsets (bottom row, 153 onsets). Locomotion onsets in both conditions activated dorsal cortex similarly. (D) As in **C**, but in an example Tlx3-Cre x Ai148 mouse that expressed GCaMP6 in L5 IT neurons during closed loop locomotion onsets (top row, 88 onsets) and open loop locomotion onsets (bottom row, 83 onsets). Note that activity decreased in posterior regions of dorsal cortex during closed loop locomotion onsets. (E) Example crystal skull craniotomy marking the 6 regions of interest in each hemisphere we selected: primary visual cortex (V1, red), retrosplenial cortex (RSC, blue), antero-medial secondary visual cortex (V2am, green), primary motor cortex (M1, yellow), anterior cingulate cortex (A24b, purple), and secondary motor cortex (M2, cyan). The white cross marks bregma. (F) Average response during closed loop locomotion onsets in C57BL/6 mice that expressed GCaMP brain wide in the 6 regions of interest (activity was averaged across corresponding regions in both hemispheres). Mean (lines) and 90% confidence interval (shading) are calculated as a hierarchical bootstrap estimate for each time bin (see Methods and **Table S1**). (G) As in **F**, but for open loop locomotion onsets. (H) As in **F**, but for visual flow onsets in the open loop condition restricted to times when the mice were not locomoting. (I) Average response during closed loop locomotion onsets in Tlx3-Cre x Ai148 mice that expressed GCaMP6 in L5 IT neurons, activity was averaged across corresponding regions in both hemispheres). Mean (lines) and 90% confidence interval (shading) are calculated as a hierarchical bootstrap estimate for each time bin (see Methods and **Table S1**). (J) As in **I**, but for open loop locomotion onsets. (K) As in **J**, but for visual flow onsets during open loop sessions restricted to times when the mice were not locomoting. (L) Similarity of the average closed and open loop locomotion onset responses quantified as the correlation coefficient between the two in a window -5 s to +3 s around locomotion onset (see Methods). Error bars indicate SEM over the 12 (6 per hemisphere) cortical regions. Statistical comparisons are against the Tlx3 data and were corrected for family-wise error rate (see Methods and **Table S1**): adjusted significance thresholds, n.s.: not significant, *: p < 0.05/9, **: p < 0.01/9, ***: p < 0.001/9.

To test whether there are differences between closed and open loop locomotion onset responses, we first compared these in mice that expressed GCaMP brain wide (C57BL/6). Consistent with previous findings (Musall et al., 2019; Stringer et al., 2019), we found that locomotion onset resulted in broad activation across most of dorsal cortex in both conditions (**Figure 1C**). However, comparing locomotion onsets in mice that expressed GCaMP6 only in Tlx3 positive L5 IT neurons, we found that the activation pattern was strikingly different between closed and open loop conditions (**Figure 1D**). While locomotion onset in the open loop condition resulted in an increase in activity across dorsal cortex, similar to that observed in C57BL/6 mice with brain wide GCaMP expression, locomotion onset in the closed loop condition resulted in a very different activation pattern: In posterior parts of dorsal cortex, the activity decreased, while in anterior parts of the dorsal cortex, the activity increased. To quantify these activation patterns across mice, we selected six regions of interest: primary visual cortex (V1), antero-medial secondary visual cortex (V2am), retrosplenial cortex (RSC), primary motor cortex (M1), anterior cingulate cortex at the level of bregma (A24b), and secondary motor cortex (M2) (**Figure 1E**). In C57BL/6 mice that expressed GCaMP brain wide, closed loop locomotion onset resulted in similar activity patterns across these six regions (**Figure 1F**). This activity pattern was not markedly different from that observed during locomotion onsets in an open loop condition (**Figure 1G**). To quantify the contribution of visual flow onset to the closed loop locomotion onset activity, we measured responses to visual flow onsets that occurred in the absence of locomotion in the open loop condition (**Figure 1H**). In C57BL/6 mice with brain wide GCaMP expression, visual flow onsets resulted in responses that were small compared to those observed during locomotion onset, even in V1. Note that a visual flow onset is a relatively weak visual stimulus compared to a drifting grating stimulus. A grating stimulus is typically composed of a gray screen or a standing grating stimulus that suddenly transitions to a drifting grating, and thus the visual flow undergoes a very high acceleration. A visual flow onset in an open loop condition is the replay of previously self-generated visual flow and thus exhibits much lower visual flow acceleration. The absence of a strong visual flow onset response in C57BL/6 mice that expressed GCaMP brain wide is consistent with a high similarity of closed and open loop locomotion onset responses.

Different from what we found in mice that expressed GCaMP brain wide, in mice that expressed GCaMP6 in Tlx3 positive L5 IT neurons, the closed loop locomotion onset patterns differed markedly when compared across the six regions of interest (**Figure 1I**). Moreover, these activation patterns were different from those observed during open loop locomotion onsets (**Figure 1J**). While these responses could partially be explained by hemodynamic occlusion, the time course of hemodynamic occlusion in Tlx3-Cre mice that expressed eGFP in L5 IT neurons was similar to that observed in C57BL/6 mice that expressed eGFP brain wide and mainly resulted in a decrease of fluorescence (**Figures S1C and S1D**). In Tlx3 positive L5 IT neurons, open loop visual flow onsets that occurred in the absence of locomotion resulted in quantifiable responses in V1 and V2am (**Figure 1K**), but the closed loop locomotion onset responses in V1 could not be explained by the sum of locomotion and visual flow onset responses (**Figures S2A and S2C**). Most of the widefield signals we record here likely originate in compartments in superficial layers which are composed of axons and dendrites of local neurons, and long-range inputs (Allen et al., 2017) (see Discussion). To test how well the widefield signals recorded in V1 reflected local somatic activity in V1, we repeated the experiments using two-photon calcium imaging of somatic activity in Tlx3 positive L5 IT neurons. We found that while there were differences between the responses measured with widefield imaging and those measured at the soma, both for locomotion onset responses and visual responses, these differences did not change our conclusions (**Figure S3**).

To quantify the differences between closed and open loop locomotion onset responses of L5 IT neurons in relation to those observed in other cortical neuron types, we repeated the experiments for the selection of excitatory and inhibitory cortical neuron types (**Figures S4A-S4I**). For each neuron type, we quantified the similarity of the activity patterns associated with closed and open loop locomotion onsets by calculating the correlation coefficient between the activity traces in a window around locomotion onset (-5 s to +3 s) for each region of interest (see Methods).

Quantifying the differences in V1, we find that the similarity in Tlx3 positive L5 IT neurons was lower than that observed in all other neuron types tested, except for Ntsr1 positive L6 neurons (**Figure S4J**). Tlx3 positive L5 IT neurons and Ntsr1 positive L6 neurons also exhibit a lower similarity than other neuron types when comparing the similarity across all regions of interest (**Figure 1L**). Both in V1 across mice, as well as when comparing across all areas, we found the highest similarity between closed and open loop locomotion onsets across regions in C57BL/6 mice that expressed GCaMP brain wide. Thus, among the neuron types tested, excitatory neurons of deep cortical layers exhibited the largest differences between closed and open loop locomotion related activation across all the selected cortical regions. The remarkable aspect of these findings is not that there are differences between closed and open loop locomotion onsets, particularly in visual regions of cortex one would perhaps expect to find differences, but that these differences are larger specifically in the deep cortical layers.

To quantify how the interaction between locomotion and visual flow feedback changes across dorsal cortex, we computed correlations between calcium activity and locomotion speed for each region of interest, for closed and open loop conditions separately. In C57BL/6 mice that expressed GCaMP brain wide, this correlation was comparable between closed and open loop conditions, was highest in posterior regions of dorsal cortex, and lower in more anterior regions (**Figures 2A and 2B**). In mice that expressed GCaMP6 in Tlx3 positive L5 IT neurons, the correlation between locomotion speed and calcium activity in the closed loop condition was negative in posterior regions of dorsal cortex and positive in anterior regions. Conversely, in open loop conditions, the correlation was positive throughout dorsal cortex (**Figures 2C and 2D**). To test whether it was the coupling of visual flow that resulted in a reduction of the correlation in the closed loop condition or whether the presence of an uncoupled visual stimulus resulted in an increase of correlation in the open loop condition, we also compared these correlations to those observed when mice were locomoting in relative darkness (see Methods). In C57BL/6 mice that expressed GCaMP brain wide, the correlation of calcium activity with locomotion in darkness was indistinguishable from that in closed or open loop (**Figure 2B**). In mice that expressed GCaMP6 in Tlx3 positive L5 IT neurons, an interesting pattern emerged. In visual regions of dorsal cortex, the correlation between locomotion speed and calcium activity in relative darkness resembled that observed in open loop conditions, while in frontal regions, it resembled that observed in closed loop conditions. In visual regions, the coupling of visual flow resulted in a reduction of correlation, while in frontal regions the presence of an uncoupled visual stimulus resulted in an increase of correlations. Thus, the difference between closed and open loop conditions likely had different origins in visual and frontal regions of dorsal cortex. Overall, we concluded that the coupling of locomotion and resulting visual flow feedback not only activated Tlx3 positive L5 IT neurons differentially but did so throughout dorsal cortex, well beyond visual regions.

**Figure 2.**
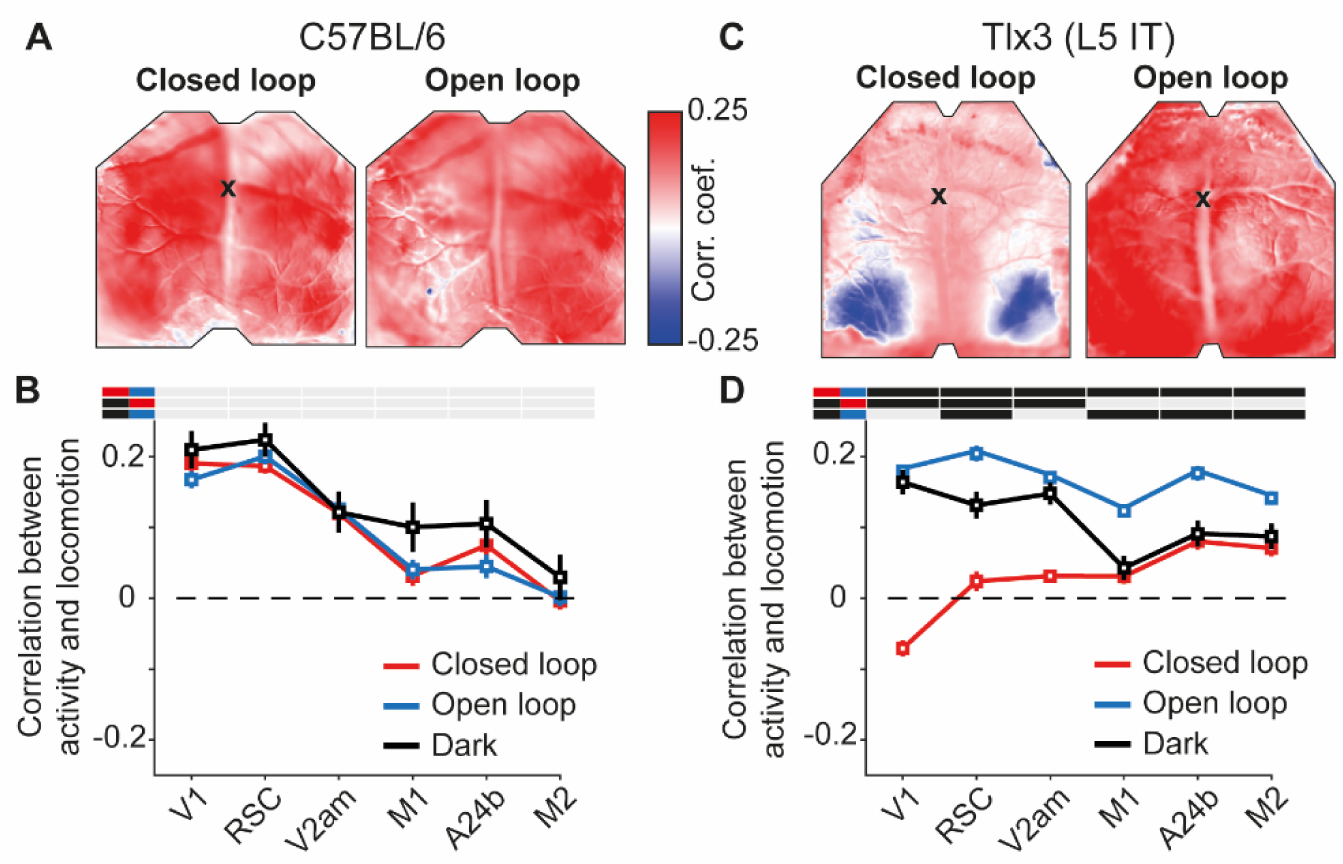
L5 IT neurons were differentially activated by locomotion depending on the type of visual feedback. (A) Correlation between calcium activity and locomotion speed in the closed loop (left) and open loop (right) conditions, calculated for each pixel in the image for one example C57BL/6 mouse that expressed GCaMP brain wide. The black cross marks bregma. Activity in most of dorsal cortex correlates positively with locomotion speed in both closed and open loop conditions. (B) Average correlation between calcium activity and locomotion speed in the 6 regions of interest in closed loop (red), open loop (blue) or dark (black) conditions in 6 C57BL/6 mice that expressed GCaMP brain wide (308, 316, and 68 5-minute recording sessions, respectively). Error bars indicate SEM over recording sessions. Bars above the plot indicate significant differences between conditions (compared conditions are indicated by colored line segments to the left, black: p < 0.05, gray: n.s., see **Table S1** for all information on statistical testing). On average, the correlation was highest in posterior dorsal cortex and was not different between conditions. (C) As in **A**, but for an example Tlx3-Cre x Ai148 mouse that expressed GCaMP6 in L5 IT neurons. Correlations of calcium activity and locomotion speed were lower in the closed loop condition compared to the open loop condition, most prominently in posterior regions of dorsal cortex. (D) As in **B**, but for 15 Tlx3-Cre x Ai148 mice that expressed GCaMP6 in L5 IT neurons (data from 420 closed loop, 394 open loop and 194 dark 5-minute recording sessions). Error bars indicate SEM over recording sessions. Correlation differed significantly between the closed and the open loop condition and was lowest in posterior dorsal cortex during the closed loop condition.

### Visuomotor prediction errors originated in visual cortical regions and had opposing effects on Tlx3 positive L5 IT activity

A consequence of the natural coupling of locomotion and visual flow feedback is that movements are a good predictor of self-generated sensory feedback. In our closed loop experiments, forward locomotion was a strong predictor of backward visual flow. It has been speculated that such predictions are used to compute prediction errors in L2/3 of cortex (Jordan and Keller, 2020).

Prediction errors come in two variants: Positive prediction errors that signal more visual flow than predicted and negative prediction errors that signal less visual flow than predicted (Keller and Mrsic-Flogel, 2018; Rao and Ballard, 1999). To probe for negative and positive prediction error signals, we quantified the responses to visuomotor mismatch events and to unexpected visual stimuli (representing less and more visual flow input than expected, respectively). Visuomotor mismatch was introduced by briefly (1 s) halting visual flow during self-generated feedback in the closed loop condition. Visual responses were elicited by presenting full-field (covering the entire toroidal screen) drifting grating stimuli to a passively observing mouse (see Methods). Please note, both types of prediction errors can be probed using a variety of stimuli, but the two we are using here, in our experience, are the most robust at eliciting strong responses in V1. In C57BL/6 mice that expressed GCaMP brain wide, both visuomotor mismatch and grating stimuli resulted in increases of activity that were strongest and appeared first in visual regions of dorsal cortex (**Figures 3A-3C and S2B**). In mice that expressed GCaMP6 in Tlx3 positive L5 IT neurons, visuomotor mismatch resulted in an increase of activity in V1, while grating onsets resulted in a decrease of activity in V1 (**Figures 3D-3F and S2D**). Thus, in dorsal cortex, both mismatch and drifting grating stimuli primarily activated visual cortex but resulted in opposing activation patterns in Tlx3 positive L5 IT neurons. An increase in activity to negative prediction errors and a decrease to positive prediction errors would be consistent with either a population of negative prediction error neurons or a type of internal representation neuron (Rao and Ballard, 1999). Given that L5 neurons in V1 do not exhibit the opposing influence of top-down and bottom-up inputs necessary to compute prediction errors (Jordan and Keller, 2020), we speculated that Tlx3 positive L5 IT neurons might function as a type of internal representation neuron.

**Figure 3.**
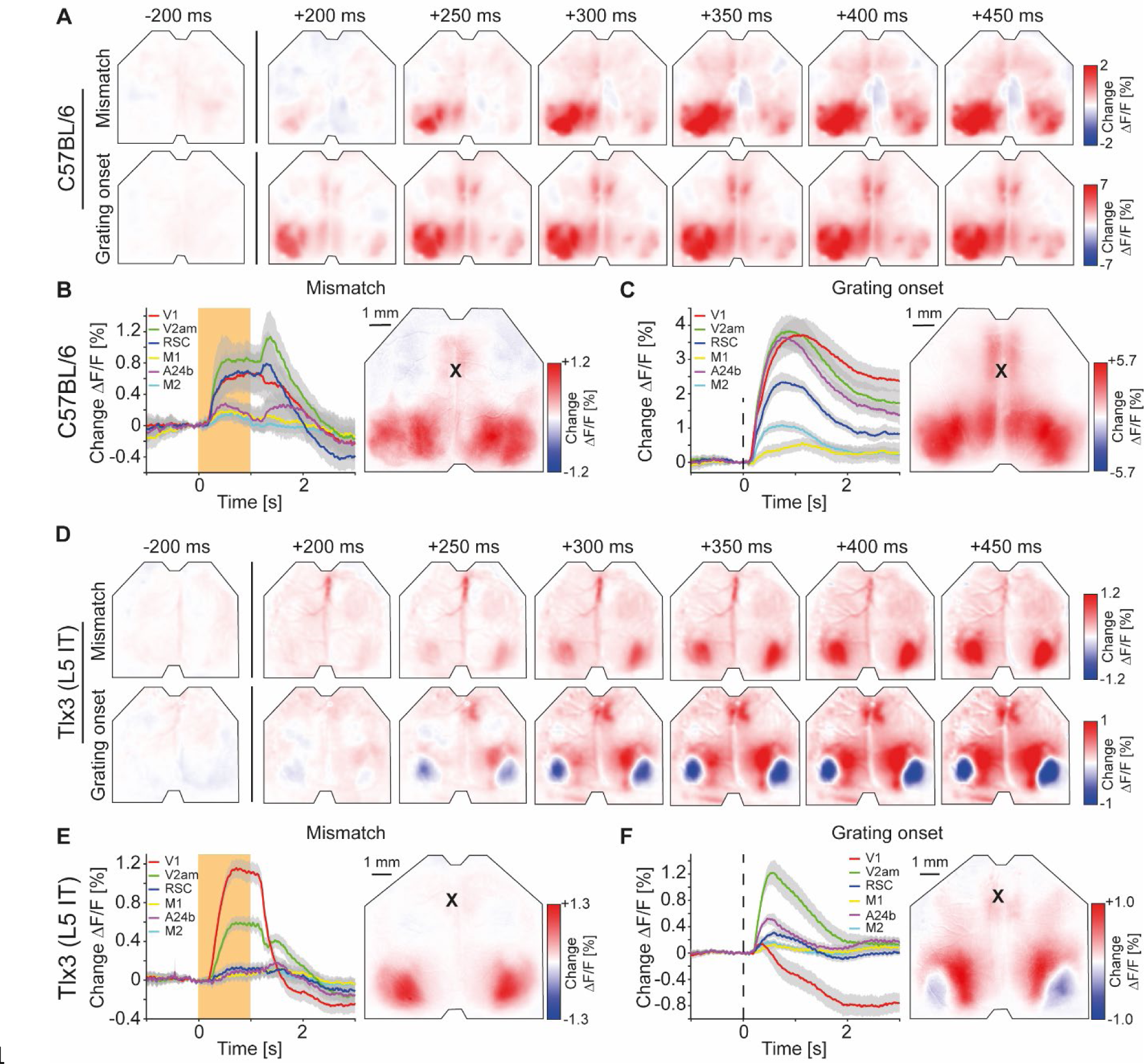
Visuomotor prediction error responses in dorsal cortex originated in V1 and activated L5 IT neurons differentially. (A) Average responses to mismatches (top row, 230 onsets) and drifting gratings (bottom row, 86 onsets) in an example C57BL/6 mouse that expressed GCaMP brain wide. (B) Left: Average response to mismatches in C57BL/6 mice that expressed GCaMP brain wide in 6 cortical regions (activity was averaged across corresponding regions in both hemispheres). Mean (lines) and 90% confidence interval (shading) are calculated as a hierarchical bootstrap estimate for each time bin (see Methods and **Table S1**). Orange shading indicates duration of mismatch event. Right: Average response map to mismatch in 6 C57BL/6 mice (see Methods). Black cross marks bregma. (C) As in **B**, but for drifting grating responses. (D) As in **A**, but in an example Tlx3-Cre x Ai148 mouse that expressed GCaMP6 in L5 IT neurons (top row: mismatch, 292 onsets; bottom row: drifting gratings, 171 onsets). (E) As in **B**, but for Tlx3-Cre x Ai148 mice that expressed GCaMP6 in L5 IT neurons. (F) As in **C**, but for Tlx3-Cre x Ai148 mice that expressed GCaMP6 in L5 IT neurons.

### The antipsychotic drug clozapine alters visuomotor integration in L5

The activation of an internal representation is the brain’s best guess at what the stimuli in the environment are. We speculated that the activation of such an internal representation would be particularly susceptible to substances that are known to reduce illusory percepts, like antipsychotic drugs. We thus quantified the changes in dorsal cortex activity associated with a single intraperitoneal injection of an antipsychotic drug (clozapine). In C57BL/6 mice that expressed GCaMP brain wide, we found that closed and open loop locomotion onsets were almost unaffected by clozapine (**Figures 4A and 4B**), while there was a small increase in open loop visual flow onset responses (**Figure 4C**). In mice that expressed GCaMP6 in Tlx3 positive L5 IT neurons, however, clozapine fundamentally changed both closed and open loop locomotion onset responses. In V1, both types of locomotion onset now resulted in a massive increase in activity (**Figures 4D and 4E**). Conversely, open loop visual flow onset responses remained largely unchanged (**Figure 4F**). These effects could not be explained by clozapine induced changes in hemodynamic occlusion (**Figure S5A**) or locomotion behavior (**Figures S5B and S5C**). Clozapine also had an opposing effect on the average correlation of activity with locomotion speed in Tlx3 positive L5 IT neurons. While the average correlation between activity and locomotion speed was increased in C57BL/6 mice that expressed GCaMP brain wide, the same measure was decreased in mice that expressed GCaMP6 in Tlx3 positive L5 IT neurons (**Figure S5D**). Finally, we confirmed that this effect was also present when recording somatic activity in V1. Using two-photon imaging, after clozapine administration we observed an increase of somatic responses during locomotion onsets that was very similar to that observed with widefield imaging (**Figures S5E and S5F**). Thus, consistent with the speculation that Tlx3 positive L5 IT neurons might function as internal representation neurons, clozapine exhibited a substantially stronger influence on locomotion onset responses in Tlx3 positive L5 IT neurons than would be expected from the effect of clozapine on brain wide responses.

**Figure 4.**
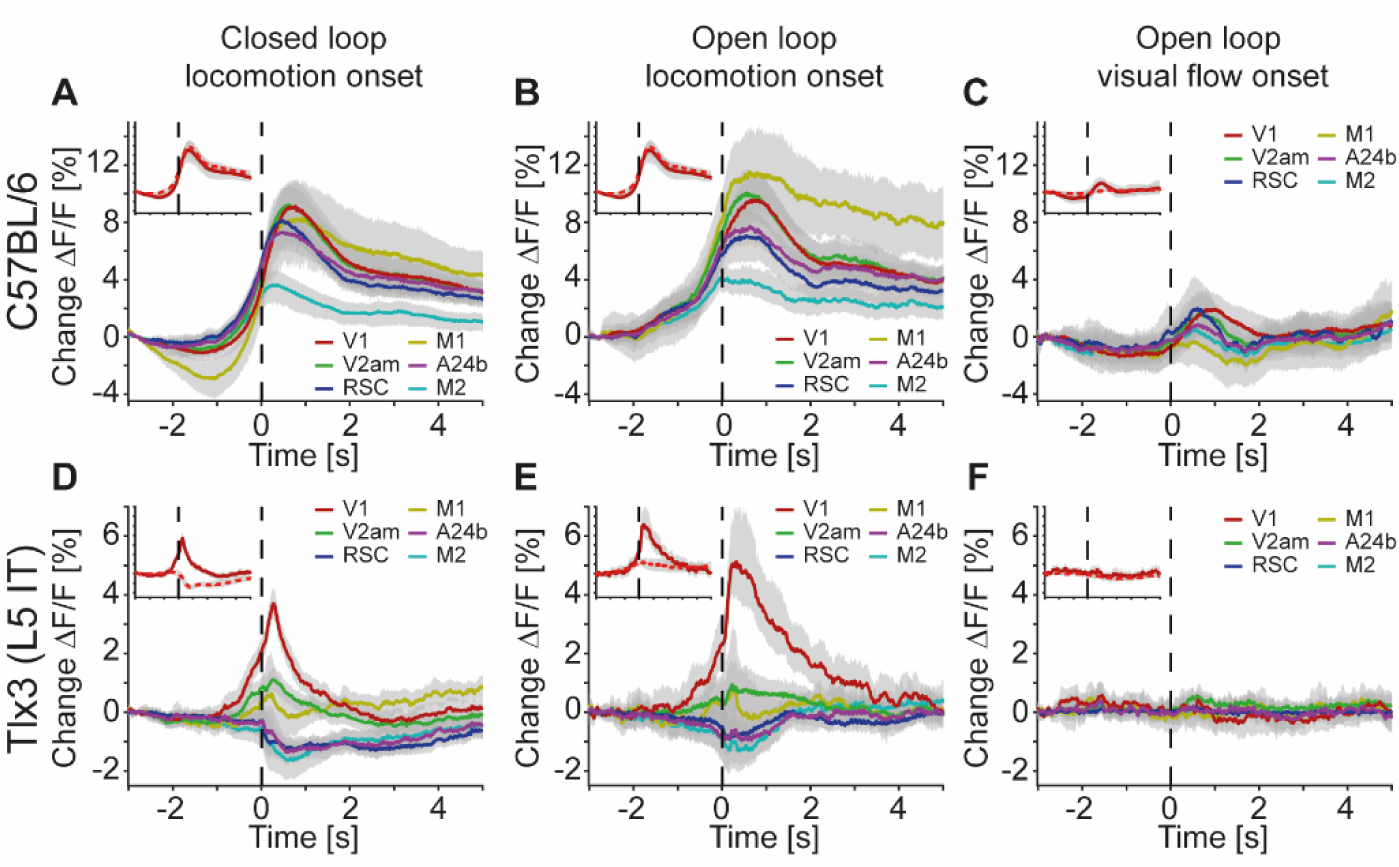
Clozapine increased locomotion related responses in L5 IT neurons in V1. (A) Average response during closed loop locomotion onsets in C57BL/6 mice that expressed GCaMP brain wide (activity was averaged across corresponding regions in both hemispheres) after a single injection of the antipsychotic drug clozapine (see Methods). Mean (lines) and 90% confidence interval (shading) are calculated as a hierarchical bootstrap estimate for each time bin (see Methods and **Table S1**). Inset: Comparison of responses for V1 in naive C57BL/6 mice (dashed red) and the same mice after clozapine injection (dark red, same data as in main panel). (B) As in **A**, but for open loop locomotion onsets after a clozapine injection. (C) As in **A**, but for open loop visual flow onsets restricted to times when the mice were not locomoting. (D) Average response during closed loop locomotion onsets in Tlx3-Cre x Ai148 mice that expressed GCaMP6 in layer L5 IT neurons (activity was averaged across corresponding regions in both hemispheres) after a single injection of the antipsychotic drug clozapine (see Methods). Mean (lines) and 90% confidence interval (shading) are calculated as a hierarchical bootstrap estimate for each time bin (see Methods and **Table S1**). Inset: Comparison of responses for V1 in naive Tlx3-Cre x Ai148 mice (dashed red) and the same mice after clozapine injection (dark red, same data as in main panel). (E) As in **D**, but for open loop locomotion onsets. (F) As in **D**, but for open loop visual flow onsets restricted to times when the mice were not locomoting.

### Antipsychotic drugs decouple long-range cortico-cortical activity

L5 IT neurons are likely the primary source of long-range cortical communication (Chen et al., 2019; Harris et al., 2019; Leinweber et al., 2017). Given that clozapine increased locomotion onset responses in L5 IT neurons in V1, we wondered whether clozapine might more generally change the long-range influence between L5 IT neurons. To investigate this possibility, we computed the correlations of calcium activity between the six regions of interest across both hemispheres and all recording conditions. We did this first in C57BL/6 mice that expressed GCaMP brain wide before (**Figure 5A**) and after a single intraperitoneal injection of clozapine (**Figure 5B**). Overall, clozapine resulted in a relatively modest decrease in correlations. On repeating this experiment in mice that expressed GCaMP6 in Tlx3 positive L5 IT neurons, we found a larger decrease in correlations that appeared to be stronger for regions that were further apart (**Figures 5C and 5D**). To quantify these changes, we plotted the correlations as a function of linear distance between the regions as measured in a top-down view of dorsal cortex (**Figure S6A**). Note, this is only an approximation of the actual axonal path length between the regions. To visualize the distribution of correlations, we interpolated the data and represented them as heatmaps (**Figure S6B**). The resulting representation for the data from C57BL/6 mice that expressed GCaMP brain wide before and after clozapine injection is shown in **Figures 6A** and **6B**, respectively. We then split the data into approximately equal portions of short- and long-range correlations using a cutoff of 0.9 times the bregma-lambda distance (approximately 3.8 mm). While both short- and long-range correlations were reduced, this reduction was not significant in either case (**Figure 6C**). We also found no evidence of a difference between short- and long-range correlation changes. In mice that expressed GCaMP6 in Tlx3 positive L5 IT neurons, however, we found a significant reduction in both short- and long-range correlations (**Figures 6D and 6E**), and both were more strongly reduced than in C57BL/6 mice that expressed GCaMP brain wide (C57BL/6 vs Tlx3-Cre x Ai148, short-range: p < 0.05; long-range: p < 10^-5^; rank-sum test). Moreover, the reduction in activity correlations was stronger for long-range correlations than it was for short-range correlations (**Figure 6F**). To control for possible clozapine induced changes in hemodynamic occlusion, we repeated the experiment and analyses in Tlx3-Cre mice that expressed eGFP in L5 IT neurons and found that alterations in hemodynamic occlusion could not account for the decrease in correlations but resulted in a small increase of the same metric (**Figures S6C-S6E**). To test whether the stress of injection, or simply passage of time, could explain the observed effects, we conducted a separate set of experiments in which we injected saline in a cohort of Tlx3-Cre x Ai148 mice and performed the same analysis. We found no evidence of a change in correlations with saline injections (**Figures S6F-S6H**). To further dissect the spatial scale at which this decorrelation occurred, we repeated the experiments in Tlx3-Cre x Ai148 mice that expressed GCaMP6 in L5 IT neurons, recording calcium activity using both a widefield and a two-photon microscope. We found that while the activity correlations measured with the widefield imaging decreased (**Figures S7A- S7C**), pairwise somatic activity correlations measured by two-photon imaging did not (**Figure S7D**).

**Figure 5.**
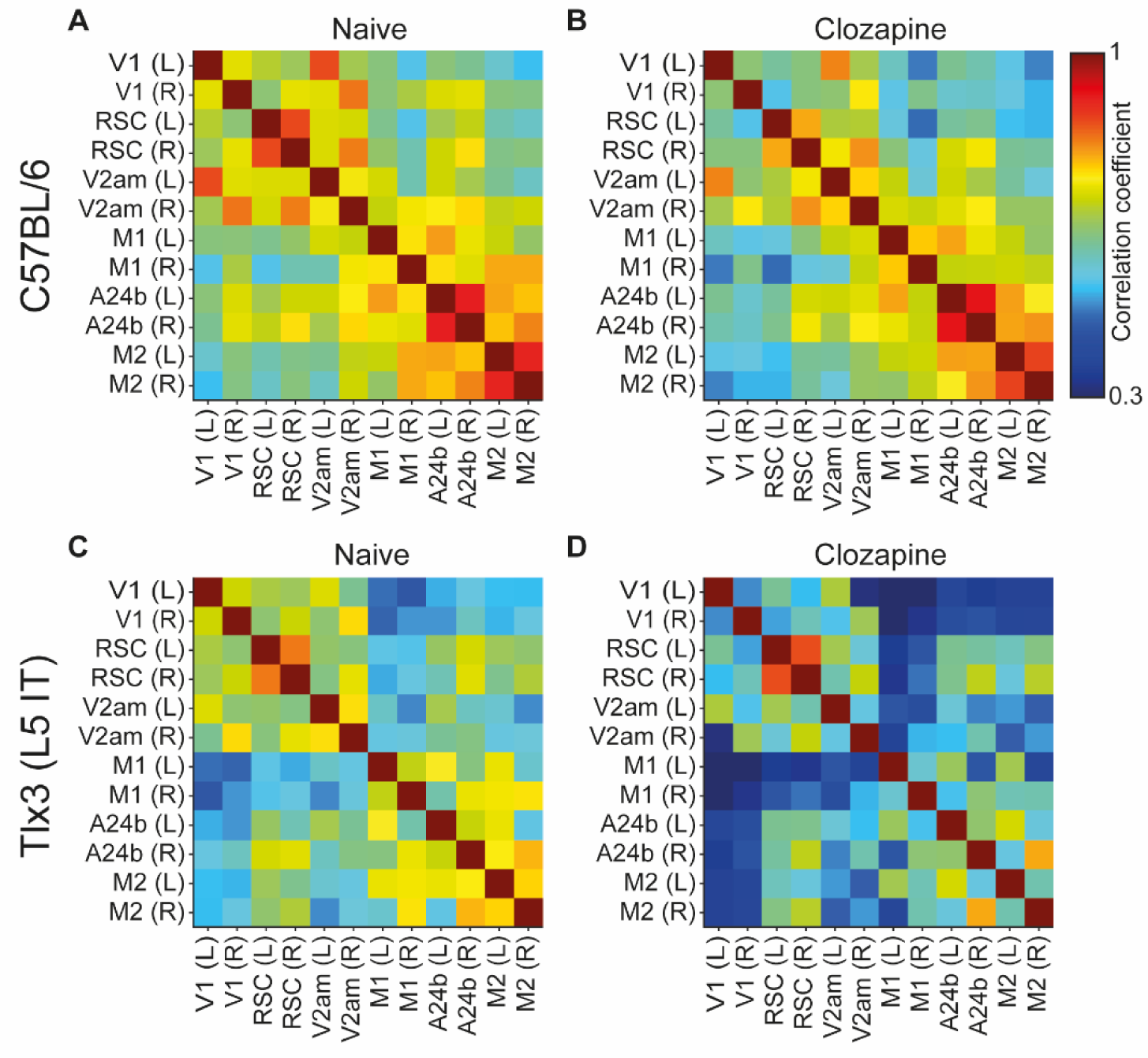
Clozapine reduced activity correlations in dorsal cortex predominantly in L5 IT neurons. (A) Average correlation of activity between the 12 regions of interest in 4 C57BL/6 mice that expressed GCaMP brain wide. (B) As in **A**, but after a single injection of the antipsychotic drug clozapine in the same 4 C57BL/6 mice. (C) Average correlation of activity between the 12 regions of interest in 5 Tlx3-Cre x Ai148 mice that expressed GCaMP6 in layer L5 IT neurons. (D) As in **C**, but after a single injection of the antipsychotic drug clozapine in the same 5 Tlx3-Cre x Ai148 mice.

**Figure 6.**
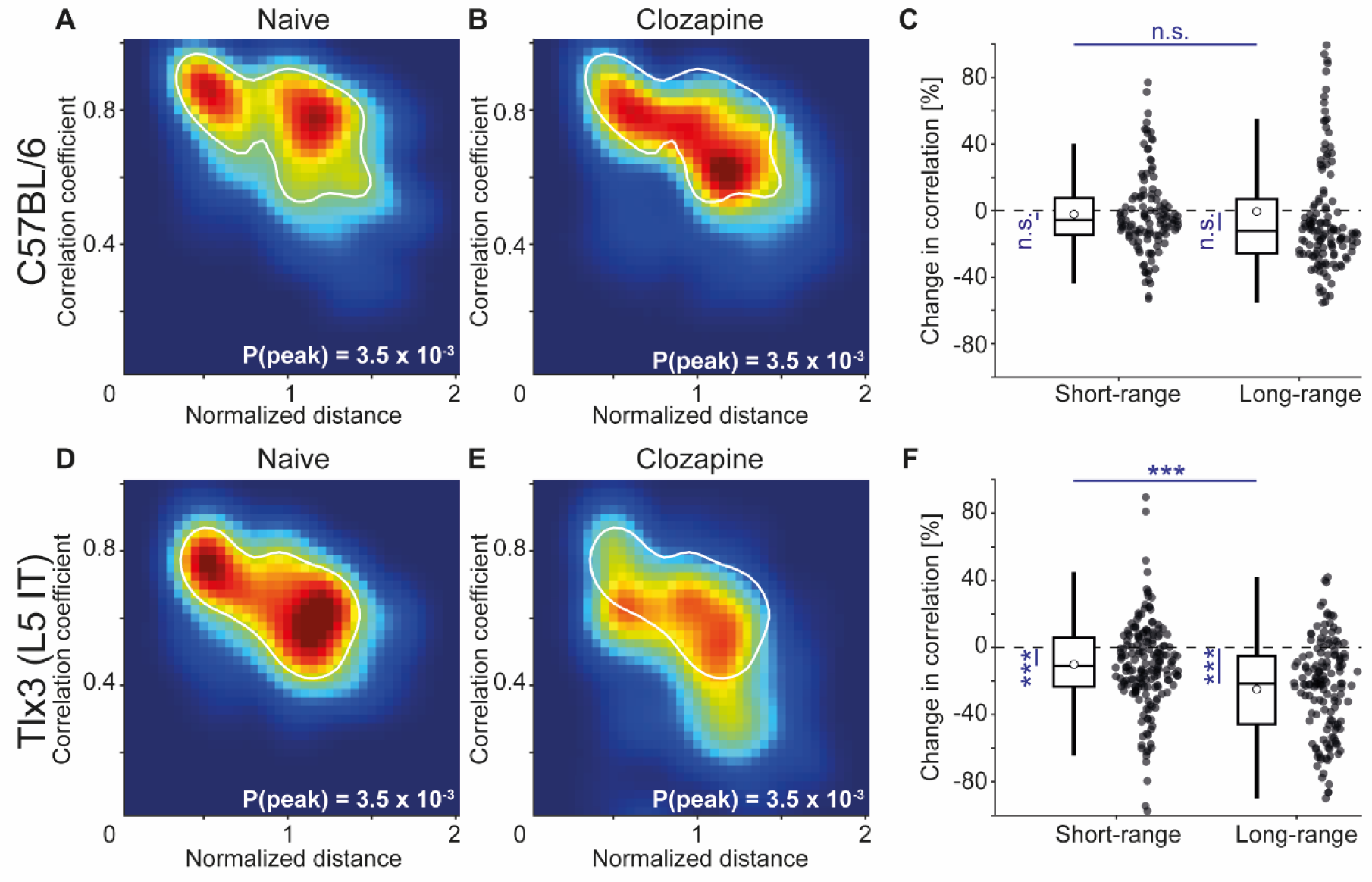
L5 IT neurons exhibited the strongest clozapine induced reduction of long-range correlations. (A) Density map of correlation coefficients as a function of distance between the regions, normalized across mice by the bregma-lambda distance (see **Figure S6A and S6B**, and Methods), for 4 naive C57BL/6 mice that expressed GCaMP brain wide. Peak density is indicated at the bottom right of the plot and corresponds to the maximum value of the color scale (dark red). The white line is a contour line drawn at 50% of peak value. (B) As in **A**, but for the same 4 C57BL/6 mice after a single injection of the antipsychotic drug clozapine. The white line is the contour line from panel **A** for comparison. (C) Clozapine induced change in the activity correlation between regions, normalized to the correlation coefficient in the naive state, in 4 C57BL/6 mice. Data were split into short- and long- range activity correlations (see Methods). Boxes represent the median and the interquartile range. The open circle indicates the mean of the distribution. The whiskers mark 1.5 times the interquartile range. Dots are the individual data points (short-range: 122 pairs of regions, long-range: 142 pairs of regions). n.s.: not significant. See **Table S1** for all information on statistical testing. (D) As in **A**, but for 5 Tlx3-Cre x Ai148 mice that expressed GCaMP6 in L5 IT neurons. (E) As in **B**, but for 5 Tlx3-Cre x Ai148 mice that expressed GCaMP6 in L5 IT neurons. The white line is the contour line from panel **D** for comparison. (F) As in **C**, but for 5 Tlx3-Cre x Ai148 mice that expressed GCaMP6 in L5 IT neurons (short-range: 182 pairs of regions, long-range: 148 pairs of regions). ***: p < 0.001.

This is consistent with the idea that decorrelation primarily occurs for long-range interactions. Finally, to confirm that the clozapine induced reduction in correlation was stronger in deep cortical layers, we repeated the experiment in a population of L2/3 and L4 excitatory neurons using Cux2- CreERT2 x Ai148 mice. We found that clozapine also decreased correlations of cortical activity in superficial excitatory neurons (**Figure S8**). However, this reduction was significantly weaker than the reduction we observed in mice that expressed GCaMP6 in Tlx3 positive L5 IT neurons (Cux2-CreERT2 x Ai148 vs Tlx3-Cre x Ai148, short-range: p < 0.005, long-range: p < 10^-8^, rank-sum test). Thus, administration of the antipsychotic drug clozapine resulted in a decorrelation of activity across dorsal cortex that was stronger in L5 IT neurons (Tlx3) than it was in either upper layer excitatory neurons (Cux2) or in the brain wide average (C57BL/6).

To test whether the decorrelation of the activity of L5 IT neurons is specific to clozapine or might more generally be a functional signature of antipsychotic drugs, we repeated the experiments with two additional antipsychotic drugs, aripiprazole and haloperidol. We found that the principal effect of decorrelation of the activity patterns of Tlx3 positive L5 IT neurons was preserved with both drugs (**Figures 7A-7F**). These changes in correlation were absent upon injection of a psychostimulant, amphetamine (**Figures 7G-7I**). Thus, it is possible that a decrease in long-range correlations between cortical L5 IT neurons might be a common mechanism of action for antipsychotic drugs.

**Figure 7.**
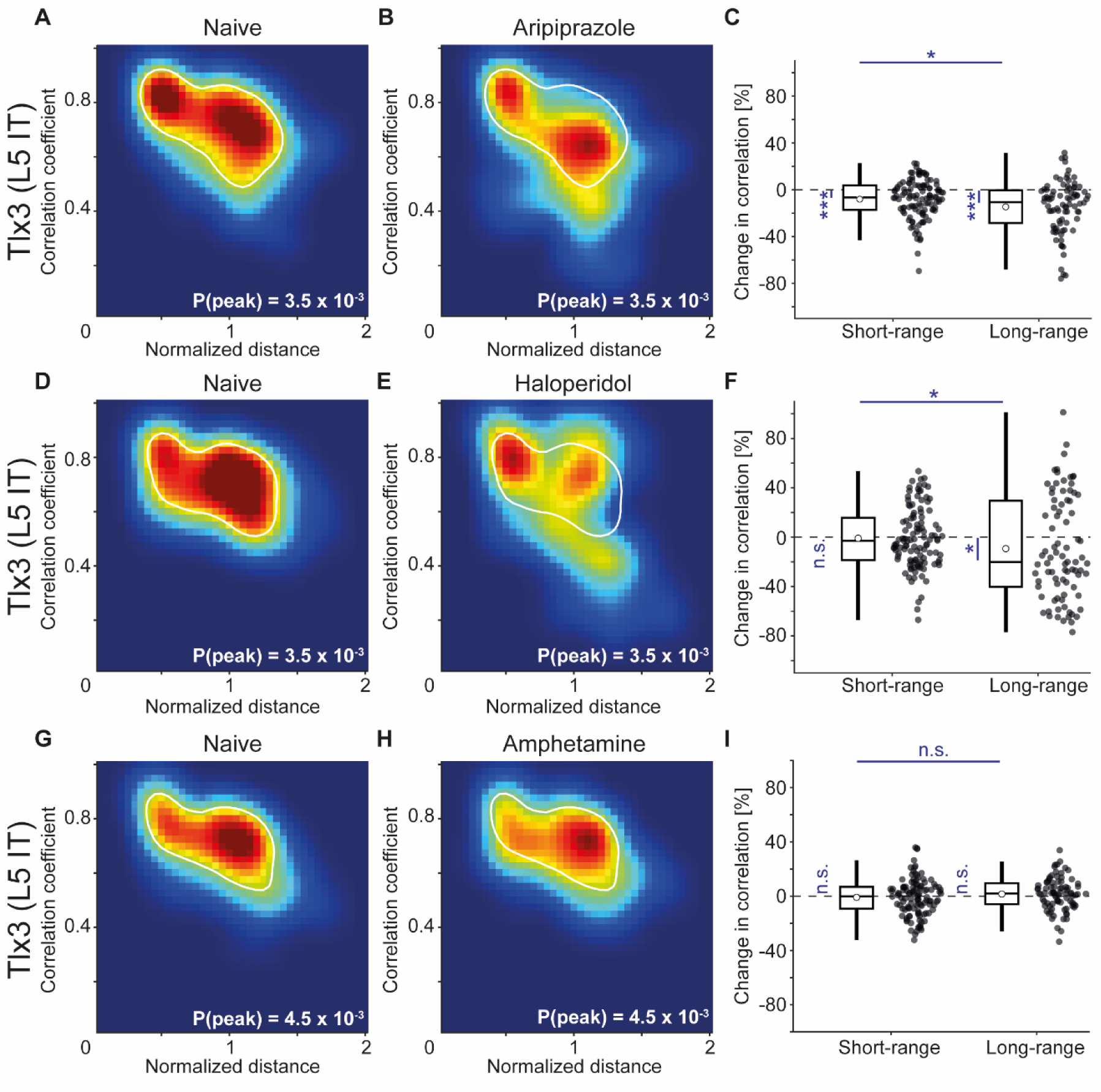
Antipsychotic drugs aripiprazole and haloperidol mimicked the decorrelation of L5 IT activity observed with clozapine while psychostimulant amphetamine did not. (A) Density map of correlation coefficients as a function of distance between the regions, normalized across mice by the bregma-lambda distance (see **Figure S6A and S6B**, and Methods), for 3 Tlx3-Cre x Ai148 mice that expressed GCaMP6 in L5 IT neurons. Peak density is indicated at the bottom right of the plot and corresponds to the maximum value of the color scale (dark red). The white line is a contour line drawn at 50% of peak value. (B) As in **A**, for the same 3 Tlx3-Cre x Ai148 mice, but after a single injection of the antipsychotic drug aripiprazole. The white line is the contour line from panel **A** for comparison. (C) Aripiprazole induced change in the activity correlation between regions, normalized to the correlation coefficient in the naive state, in 3 Tlx3-Cre x Ai148 mice. Data were split into short- and long-range activity correlations (see Methods). Boxes represent the median and the interquartile range. The open circle indicates the mean of the distribution. The whiskers mark 1.5 times the interquartile range. Dots are the individual data points (short-range: 114 pairs of regions, long-range: 84 pairs of regions). *: p < 0.05, ***: p < 0.001. See **Table S1** for all information on statistical testing. (D) As in **A**, but for 3 different Tlx3-Cre x Ai148 mice. (E) As in **D**, for the same 3 Tlx3-Cre x Ai148 mice, but after a single injection of the antipsychotic drug haloperidol. The white line is the contour line from panel **D** for comparison. (F) As in **C**, but for the 3 mice that had received a single injection of the antipsychotic drug haloperidol (short-range: 114 pairs of regions, long-range: 84 pairs of regions). *: p < 0.05; n.s.: not significant. (G) As in **A**, but for 3 different Tlx3-Cre x Ai148 mice. (H) As in **G**, for the same 3 Tlx3-Cre x Ai148 mice, but after a single injection of the psychostimulant amphetamine. The white line is the contour line from panel **G** for comparison. (I) As in **D**, but for the 3 mice that had received a single injection of the psychostimulant amphetamine (short-range: 114 pairs of regions, long-range: 84 pairs of regions). n.s.: not significant.

Changes in the correlation between cortical regions could either be driven by changes in correlated external inputs or through changes in communication between cortical regions. If antipsychotic drugs primarily reduce correlations by reducing the strength of communication between cortical regions, independent of whether this communication is direct or indirect, we would expect to find changes in the way responses spread between cortical regions. We used visuomotor prediction error responses that originated in V1 (**Figure 3**) to quantify how the spread of these responses in Tlx3 positive L5 IT neurons was influenced by antipsychotic drugs. The computation of negative prediction errors during movement is thought to be driven by an excitatory prediction of visual flow mediated by long-range cortical input from L5 neurons in regions like A24b (Leinweber et al., 2017), as well as an inhibitory bottom-up visual input. By contrast, the computation of a positive prediction error during passive observation is thought to be driven by an excitatory bottom-up visual input and an inhibitory top-down input. Thus, we would primarily expect to find a reduction in mismatch responses in L5 IT neurons and a reduction in the spread of this signal to secondary visual regions.

This is indeed what we observed: Mismatch responses were partially reduced in V1 and almost absent in V2am after injection of antipsychotic drugs (**Figure 8A**). While responses to onsets of drifting gratings were also reduced in V2am, they were overall less affected by the antipsychotic drugs than mismatch responses (**Figure 8B**). These results are consistent with the interpretation that antipsychotic drugs reduce lateral communication in L5. Whether this communication is mediated by direct long-range connections between L5 neurons, or indirectly via other cortical neurons or subcortical loops, remains to be tested.

**Figure 8.**
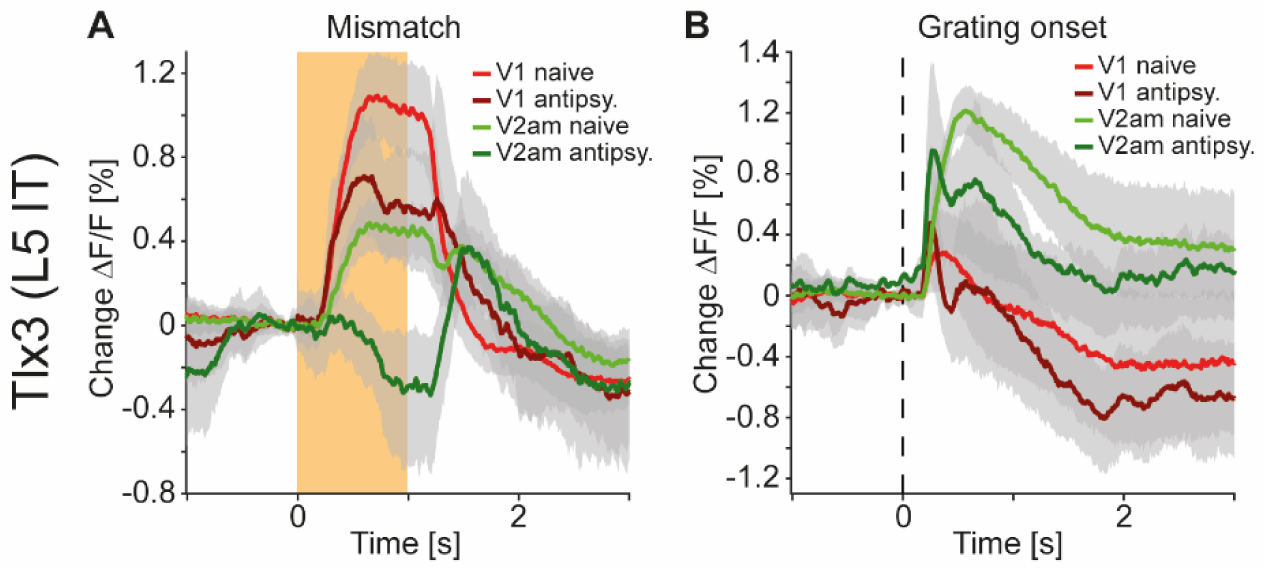
Antipsychotic drug treatment preferentially reduced responses to and propagation of negative prediction errors. (A) Average responses to mismatches in Tlx3-Cre x Ai148 mice that expressed GCaMP6 in L5 IT neurons before (red: V1 naive, green: V2am naive, activity was averaged across corresponding regions in both hemispheres) and after (dark red: V1, dark green: V2am) injection of a single dose of an antipsychotic drug (data were averaged over all antipsychotics used: Clozapine: 5 mice, aripiprazole: 3 mice, and haloperidol: 3 mice). Orange shading indicates duration of the mismatch event. Mean (lines) and 90% confidence interval (shading) are calculated as a hierarchical bootstrap estimate for each time bin (see Methods and **Table S1**). (B) As in **A**, but for drifting grating onsets.

## DISCUSSION

When interpreting our results, it should be kept in mind that the source of the widefield imaging signal in these experiments is not entirely clear and is likely a combination of somatic, dendritic, and axonal signals. Given that the signal attenuation half depth in cortex is on the order of 100 µm (Aravanis et al., 2007), most signals likely come from the superficial 100 µm of tissue (Allen et al., 2017). In the case of the Tlx3 recordings, this would mean that we were recording a mixture of axonal and dendritic signals of the Tlx3 neurons that have their soma in L5 but have a dense net of axons and dendrites in L1. In the case of the brain wide expression of GCaMP using an AAV-PHP.eB in C57BL/6 mice, the signals we recorded likely also include contributions from thalamic axons. In addition to this, there is a hemodynamic contribution to the signal (Allen et al., 2017; Ma et al., 2016; Valley et al., 2020) that is the result of varying occlusion by blood vessels. We attempted to correct for the hemodynamic response using a 405 nm wavelength isosbestic illumination of GCaMP6 but could not recover the hemodynamic responses measured in eGFP experiments.

Accurate correction likely requires calibration on eGFP measurements (Valley et al., 2020). Given that hemodynamic responses should not systematically differ between different mouse lines, and that in the crystal skull preparation the contribution of hemodynamic occlusion to increases in fluorescence was relatively small (**Figure S1**), these hemodynamic artefacts should not substantially alter our conclusions. Lastly, it should be kept in mind that responses in these widefield recordings inherently reflect population averages. Assuming there is some form of lateral inhibition among neurons, population averages need not reflect responses of neurons that are tuned to the specific stimulus presented. For example, the suppression to grating stimuli we observed in V1 Tlx3 positive L5 IT neurons could be the result of a subset of Tlx3 neurons that are strongly responsive to the specific grating presented suppressing the rest of the population of Tlx3 neurons. The sum of both could then result in a net decrease in calcium activity. Thus, an understanding of the mechanism that results in opposing population responses of Tlx3 positive L5 IT neurons to positive and negative prediction errors will require measurements with single neuron resolution.

In our experiments we have focused on Tlx3 positive L5 IT neurons, but given that L5 IT neurons are densely connected to L5 pyramidal tract neurons (Anderson et al., 2010; Kiritani et al., 2012) and to L6 neurons (Xu et al., 2016; Yamawaki and Shepherd, 2015), it is not unlikely that these neuron types would follow similar patterns of activity and exhibit similar susceptibility to the influence of antipsychotic drugs. In the comparison of closed and open loop locomotion onsets, we do indeed find that Ntsr1 positive L6 excitatory neurons exhibit similar levels of dissimilarity as those observed in Tlx3 positive L5 IT neurons (**Figure 1L**). It is possible that responses in the Ntsr1 recordings, however, were noisier than those in the Tlx3 recordings (**Figure S4**), likely because of the lower dendritic and axonal volume of Ntsr1 positive neurons in L1. Thus, while we have focused on the Tlx3 positive L5 IT neurons mainly for reasons of experimental convenience, it is possible that both the differential activation in closed and open loop locomotion onsets as well as the susceptibility of long-range correlations to antipsychotic drugs is a general feature of cortical layers 5 and 6. Another possibility we currently cannot entirely rule out is that the decorrelation effect we observed in Tlx3 positive L5 IT neurons is specific to the Tlx3-Cre mouse line. However, this seems unlikely given that all mouse lines we used were maintained on the same C57BL/6 background.

It is still unclear what the relevant sites of action in the brain are of antipsychotic drugs. Antipsychotic drugs are often antagonists of the D2 receptors, which are most strongly expressed in striatum. Consequently, it is sometimes assumed that the relevant site of action of antipsychotics is the striatum. Consistent with this, an increase in striatal dopamine activates auditory cortex (Maia and Frank, 2017) and mediates hallucination-like perceptions in mice (Schmack et al., 2021). It is intriguing, however, to speculate that the effects we observed in L5 IT neurons, which provide a dominant input to striatum (Morita et al., 2019), may be an alternate site of action. While it is possible that the primary site of action of the antipsychotic drugs with respect to the L5 decorrelation phenotype we observed is striatum, this is unlikely. Changes to striato-thalamo- cortical loops could appear as changes in functional connectivity observed in cortex. However, feedback from striatum to cortex is exclusively mediated through thalamus, and neither L4, the primary thalamo-recipient layer in cortex, nor our brain wide recordings exhibited any of the effects we saw in L5 IT neurons. Thus, the effects we saw in L5 IT neurons are either not inherited from striatum or are mediated by a separate set of thalamic neurons via direct thalamic input to deep cortical layers (Constantinople and Bruno, 2013; Douglas and Martin, 1991). Thalamocortical collaterals in deep layers, however, come from thalamic neurons that project primarily to superficial layers (Oberlaender et al., 2012). Thus, we think it is unlikely that the observed effects in L5 IT neurons were inherited from striatum. Independent of whether the treatment-relevant site of action of antipsychotic drugs is indeed in L5 IT neurons, or whether the observed decorrelation of activity is relevant to antipsychotic potency of these drugs, we speculate that the long-range decorrelation of activity could be used as a basis for a functional screen for compounds with antipsychotic efficacy.

Given that in the brain wide recordings the decorrelation of activity was weaker compared to that observed in Tlx3 positive L5 IT neurons, it might be possible to test for similar effects in human subjects using layer specific fMRI recordings (Haarsma et al., 2020).

A rather surprising implication of our results is that the dominant mode of coupling in cortex might be lateral, not vertical. Two findings would argue for this interpretation. First, while L5 was differentially activated across cortical regions during closed and open loop locomotion onsets, L2/3 exhibited rather limited differences. Second, antipsychotics decorrelated L5 activity but had no effect on L2/3. Both findings could be explained if the long-range lateral coupling within L5 is stronger than the local vertical coupling between L2/3 and L5.

One of the diseases treated with antipsychotic drugs is schizophrenia. It has been speculated that disrupted resting-state functional connectivity in brain networks involving cortex is central to the pathogenesis of schizophrenia (Li et al., 2017; Northoff and Duncan, 2016). Treatment-naïve schizophrenia patients predominantly exhibit increases in resting state functional connectivity between cortical regions, while medicated schizophrenia patients exhibit more balanced fractions of increased and decreased functional connections (Anticevic et al., 2015; Du et al., 2021; Li et al., 2017). Auditory hallucinations, one of the hallmark symptoms of schizophrenia, are associated with increased activation of diverse cortical regions (Catafau et al., 1994; Dierks et al., 1999; Shergill et al., 2000). Thus, it is conceivable that antipsychotic drugs counteract this by reducing the strength of long-range cortical connectivity mediated by L5 IT neurons, independent of whether the increased activity in schizophrenic patients is indeed driven by aberrant increases in the strength of long-range cortical connections or is a consequence of an aberrant increase in the strength of striato-thalamo- cortical loops. However, a direct involvement of cortical circuits in the pathogenesis of schizophrenia is consistent with the glutamate hypothesis of schizophrenia. This hypothesis is based on the finding that NMDA receptor antagonists induce schizophrenia-like symptoms (Moghaddam and Javitt, 2012), and that disturbances of NMDA and AMPA receptor related gene expression are associated with schizophrenia (Harrison and Weinberger, 2005; Singh et al., 2020). In V1, it has been shown that NMDA receptors are necessary during visual development to establish normal prediction error signaling (Widmer and Keller, 2021). Thus, it is conceivable that a disruption of glutamatergic signaling in cortex that alters the way internal representations are activated by prediction errors is central to the etiology of schizophrenia. If so, we should find impairments in all domains of cortical function, including sensory processing. In the case of the sensory cortex, cortical dysfunctions should be apparent earlier in development, concurrent with the critical period for plasticity in sensory areas of cortex. While the core symptoms of schizophrenia typically appear in late adolescence, sensory processing is indeed impaired in schizophrenic patients (Javitt, 2009) and is apparent, for example, in the reduction of surround suppression in V1 (Yoon et al., 2009). Interestingly, some of these disturbances are detectable already during childhood as visuo-perceptual and reading anomalies in some subjects that later develop schizophrenia (Parellada et al., 2017). Positive symptoms of schizophrenia can be explained in a predictive processing framework of cortical function as a disturbance of a prediction error based update of internal representations (Fletcher and Frith, 2009; Frith, 2005). There are two types of prediction error neurons, positive and negative, and, given that cortex likely implements a non-hierarchical variant of predictive processing (Garner and Keller, 2022), two types of internal representation neurons. One set of internal representation neurons is excited by positive prediction errors and inhibited by negative prediction errors, and the other one is inhibited by positive prediction errors and excited by negative prediction errors (Keller and Mrsic-Flogel, 2018). Tlx3 positive L5 IT neurons exhibited opposing responses to positive and negative prediction errors (**Figure 3**), and, while several different interpretations are conceivable, these responses would be consistent with those of internal representation neurons. Given that the effects of antipsychotic drugs are predominantly observed in L5 IT neurons, and that anesthesia is thought to decouple L5 neuron soma from their apical tuft and thereby prevent conscious perception (Suzuki and Larkum, 2020), we speculate that Tlx3 positive L5 IT neurons may be one type of internal representation neuron and one of the functionally relevant sites of antipsychotic action.

## METHODS

### Key Resource Table

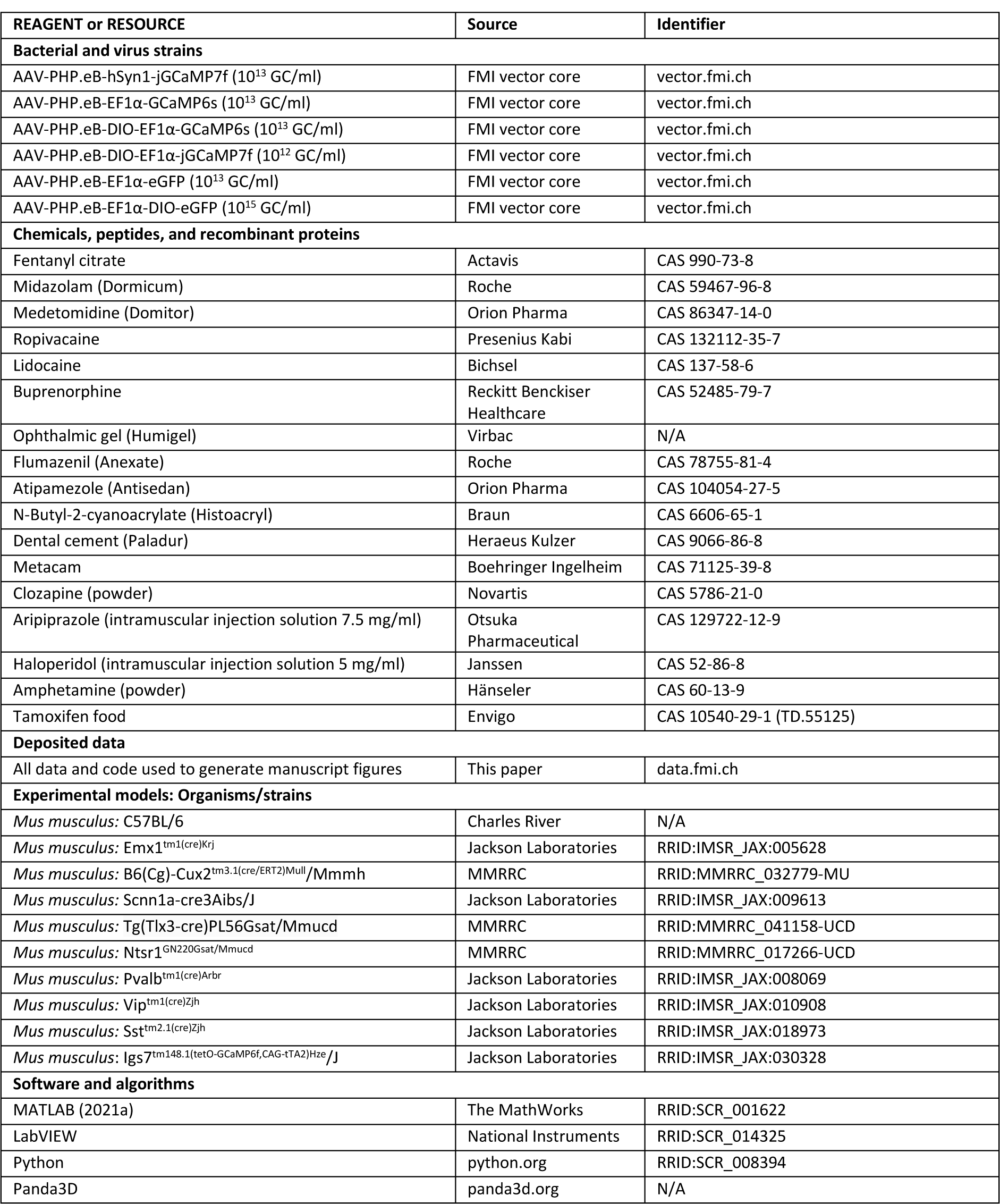

### Mice

All animal procedures were approved by and carried out in accordance with guidelines of the Veterinary Department of the Canton Basel-Stadt, Switzerland. The mice used for cell-type specfic targeting and subsequent widefield imaging in this study were kept on a C57BL/6 background and were of the following genotype: 6 C57BL/6 (Charles River Laboratories), 4 Emx1-Cre to target expression to cortical excitatory neurons (Gorski et al., 2002), 4 Cux2-CreERT2 to target expression to L2/3 and L4 excitatory neurons (Franco et al., 2012), 7 Scnn1a-Cre to target expression to a subset of L4 excitatory neurons (Madisen et al., 2010), 25 Tlx3-Cre to target expression to a subset of L5 IT neurons (Gerfen et al., 2013), 3 Ntsr1-Cre to target expression to a subset of L6 excitatory neurons (Gong et al., 2007), and, to target inhibitory subpopulations, we used 2 PV-Cre (Hippenmeyer et al., 2005), 6 VIP-Cre (Taniguchi et al., 2011), and 5 SST-Cre (Taniguchi et al., 2011) mice. Ai148 ((Daigle et al., 2018), Jackson Laboratories, stock #030328) mice were used as breeders to drive GCaMP6f expression in Cre positive neurons. An additional 8 C57BL/6 and 7 Tlx3-Cre mice were used in experiments controlling for hemodynamic responses. To induce expression of Cre in the Cux2- CreERT2 line, mice were provided for at least 1 week with tamoxifen containing food (Envigo, 400 mg tamoxifen per kg of food) as their sole food source. Mice were group housed in a vivarium (light/dark cycle: 12/12 hours). Experimental mice were of either sex.

### Surgery and virus injections

For all surgical procedures, mice were anesthetized using a mixture of fentanyl (0.05 mg/kg; Actavis), midazolam (5.0 mg/kg; Dormicum, Roche), and medetomidine (0.5 mg/kg; Domitor, Orion). In a subset of mice (6 C57BL/6, 4 Emx1-Cre, and 4 SST-Cre mice), we injected an AAV vector based on the PHP.eB capsid (Chan et al., 2017) retro-orbitally (6 μl each eye of at least 10^13^ GC/ml) to drive expression of GCaMP under either the EF1α or hSyn1 promoter pan-neuronally, or in a cell type specific manner using a double-floxed inverted open reading frame (DIO) construct. For hemodynamic response control experiments, we used an AAV-PHP.eB-EF1α-eGFP to pan-neuronally express eGFP (**Figures S1A and S1B)** or an AAV-PHP.eB-EF1α-DIO-eGFP to restrict eGFP expression to L5 IT neurons in Tlx3-Cre mice (**Figures S1C and S1D, S5A, S6C-S6E**). To improve optical access to the cortex, we implanted crystal skull cranial windows (Kim et al., 2016). Prior to removing the skull plate overlying dorsal cortex, we recorded the location of bregma relative to other landmarks on the skull. Superglue (Pattex) was used to glue the crystal skull in place in the craniotomy. In a subset of mice, for the comparison of hemodynamic responses (**Figure S1A and S1B**), we used a clear skull preparation (Guo et al., 2014). To do this, we attached the crystal skull coverslip directly onto the cleaned skull surface with a three-component polymer (C&B Metabond, Parkell). A custom-machined titanium head bar was attached to the skull using dental cement (Paladur, Heraeus). An epifluorescence overview image was taken to mark reference points on the dorsal cortical surface. Anesthesia was antagonized by an intraperitoneal injection of a mixture of flumazenil (0.5 mg/kg; Anexate, Roche) and atipamezole (2.5 mg/kg; Antisedan, Orion Pharma). For peri- and post-operative analgesia we injected buprenorphine (0.1 mg/kg; Reckitt Benckiser Healthcare (UK) Ltd.) and Metacam (5 mg/kg; Boehringer Ingelheim). Imaging commenced at the earliest 1 week after head bar implantation or 3 weeks after retro-orbital AAV injection. Clozapine (Novartis), aripiprazole (Otsuka Pharmaceutical), haloperidol (Janssen), amphetamine (Hänseler), or saline (0.9% NaCl in H2O) was intraperitoneally injected at 0.2 μg/g - 10 μg/g, 0.2 μg/g, 0.1 μg/g, 4 μg/g body weight, or undiluted, respectively.

### Virtual reality setup and stimulus design

For all experiments we used a virtual reality setup as previously described (Leinweber et al., 2014). Mice were head-fixed and free to run on a spherical, air-supported Styrofoam ball. A virtual corridor was projected (using a Samsung SP-F10M projector) onto a toroidal screen positioned in front of the mouse covering a field of view of approximately 240 degrees horizontally and 100 degrees vertically. Recordings were blocked into 5-minute sessions.

Experiments typically commenced with 4 sessions of a closed loop condition in which the rotation of the spherical treadmill was coupled to the movement in a virtual corridor. To introduce mismatches, we broke the coupling between locomotion and visual flow by briefly halting visual flow for 1 s at random times. Closed loop sessions were followed by 4 sessions of an open loop condition, in which rotation of the spherical treadmill and movement in the virtual corridor were decoupled. The visual stimulus consisted of a replay of the visual flow recorded in the previous closed loop condition.

Following this we recorded 2 sessions of what we refer to as a dark condition in which the virtual reality was switched off. Please note, this was not complete darkness, primarily because the shielding of the blue excitation light was not light proof. Finally, we recorded activity in 2 sessions of a grating condition in which we presented drifting grating stimuli covering the full toroidal screen placed in front of the mouse (8 directions of 6 s ± 2 s (mean ± SD) duration each with an inter-grating interval of 4.5 s ± 1.5 s (mean ± SD) gray screen, presented in randomized order). These sessions, totaling approximately 1 hour of recording time per day, were acquired for 3 consecutive days. For mice in which we went on to test the effect of drugs on dorsal cortex activity, the first post-drug injection data were acquired +1 h after drug injection (+30 mins in the case of amphetamine because of faster onset kinetics of the drug) and further data were collected at the +24 h and +48 h timepoints. Data in the manuscript are shown as the combined data over either all data collected before drug injection (labelled as naive) or after substance injection (labelled as clozapine, aripiprazole, haloperidol, amphetamine, or saline, respectively).

### Data acquisition

All widefield imaging experiments were performed on a custom-built macroscope consisting of commercially available objectives mounted face-to-face (Nikon 85 mm/f1.8 sample side, Nikon 50 mm/f1.4 sensor side). We used a 470 nm LED (Thorlabs) powered by a custom-built LED driver for exciting GCaMP (or eGFP) fluorescence through an excitation filter (SP490, Thorlabs) reflected off a dichroic mirror (LP490, Thorlabs) placed in the parfocal plane of the objectives. Green fluorescence was collected through a 525/50 emission filter on an sCMOS camera (PCO edge 4.2).

Apertures on objectives were usually kept maximally open and the current at the LED driver was used to adjust fluorescence intensity to a value that was kept below 25% of the maximum dynamic range of the sensor. In cases where this was not possible (e.g., transfection of eGFP yielded extremely bright fluorescence, often visible with the naked eye), objective apertures were gradually reduced to avoid overexposure of the sensor. LED illumination was adjusted with a collimator (Thorlabs SM2F32-A) to achieve homogenous illumination across the surface of the cranial window. The resulting Gaussian profile of the illumination cone was further trimmed with black tape on the sample side objective to avoid light shining directly into the mouse’s eye. An Arduino board (Arduino Mega 2560) was used to control LED onsets synced to the frame trigger signal of the camera. The duty cycle of the 470 nm LED was 90%. Raw images were acquired at full speed (100 Hz; except for eGFP controls, which were acquired at 50 Hz effective frame rate) and full dynamic range (16 bit) of the sensor. The raw images were cropped on-sensor and the resulting data was streamed to disk with custom written software in LabVIEW (National Instruments), resulting in an effective pixel size of 60 μm^2^ at a standardized imaging resolution of 1108 pixels x 1220 pixels (1.35 MP).

Two-photon calcium imaging was performed using a custom-built microscope based on a Thorlabs B-scope (Leinweber et al., 2014). Illumination source was a tunable femtosecond laser (Coherent Chameleon) tuned to 930 nm. Emission light was band-pass filtered using a 525/50 filter for GCaMP and detected using a GaAsP photomultiplier (H7422, Hamamatsu). Photomultiplier signals were amplified (DHPCA-100, Femto), digitized (NI5772, National Instruments) at 800 MHz, and band-pass filtered around 80 MHz using a digital Fourier-transform filter implemented in custom-written software on an FPGA (NI5772, National Instruments). The scanning system of the microscope was based on a 12 kHz resonant scanner (Cambridge Technology). Images were acquired at a resolution of 750 pixels x 400 pixels at a 60 Hz frame rate and a piezo-electric linear actuator (P-726, Physik Instrumente) was used to move the objective (Nikon 16x, 0.8 NA) in steps of 15 µm between frames to acquire images at 4 different depths. This resulted in an effective frame rate of 15 Hz. The field of view was 350 µm x 350 µm.

### Data processing

Off-line data processing and data analysis (section below) were done with custom-written MATLAB (2021a) scripts. For widefield macroscope imaging, raw movie data was manually registered across days by aligning subsequent mean projections of the data to the first recorded image sequence. We placed regions of interest (ROIs) relative to readily identifiable anatomical landmarks that had been previously noted during cranial window surgery (see above). This resulted in the selection of six 20 pixels x 20 pixels ROIs per hemisphere. Of those ROIs, we calculated the activity as the ΔF/F0, wherein F0 was the median fluorescence of the recording (approximately 30 000 frames in a 5 min recording). ΔF/F0 was corrected for slow fluorescence drift caused by thermal brightening of the LED using 8^th^ percentile filtering with a 62.5 s moving window similar to what was described previously for two-photon imaging (Daniel A. Dombeck et al., 2007).

Raw two-photon imaging data were full-frame registered to correct for brain motion. Neurons were manually selected based on mean and maximum fluorescence images. Raw fluorescence traces were corrected for slow drift in fluorescence using an 8^th^ percentile filtering with a 15 s window (Daniel A Dombeck et al., 2007). ΔF/F traces were calculated as mean fluorescence in a selected region of every imaging frame and subsequently subtracted and normalized by the median fluorescence.

### Data analysis

Locomotion and visual flow onsets were determined based on threshold crossing of locomotion or visual flow speed. To select only well isolated onsets, we used a speed threshold of 30 cm/s and excluded all onsets with locomotion or visual flow in the 3 s window preceding the onset. To increase the number of locomotion onsets that passed the speed threshold criterion for the eGFP imaging data (**Figures S1, S5A**), we excluded only those onsets if there was locomotion in the 1 s window preceding onset.

All stimulus response curves (**Figures 1F-1L, 3B and 3C, 3E and 3F, 4, 8, S1-S4, S5A and S5B, S5E and S5F**) were baseline subtracted. The baseline subtraction window for unpredictable stimuli (mismatches and grating onsets) was -200 ms to 0 ms before stimulus onset. For closed loop and open loop locomotion onsets, and open loop visual flow onsets, to accommodate calcium indicator offset dynamics and potential anticipatory activity, we placed the baseline subtraction windows for these onsets at -2900 ms to -2700 ms for GCaMP imaging, or -900 ms to -700 ms for eGFP imaging. Varying this window within the bounds of offset dynamics of the indicator and anticipatory activity onset did not change the conclusions drawn in this manuscript. Analyses in which image sequences were aligned to event onsets (**Figures 1C-1D and 3**) used the same parameters as above.

On rare occasions, we observed seizure-like activity after antipsychotic drug injection. These events resulted in atypically high levels of correlation, and to avoid them from influencing our conclusions we excluded these data from further analysis. We excluded sessions if they contained an event in which average activity of all 12 ROIs remained elevated above 30% ΔF/F for more than 10 s. This criterion led to the exclusion of 0.26% of data (10 out of 3801 of 5 min recording sessions).

To quantify the similarity of closed loop and open loop locomotion onset responses shown in **Figure 1L** we calculated the correlation coefficient between the averaged responses (onsets and mice, 1 s moving average) in a window -5 s to +3 s, for each ROI. For **Figure S4J**, we first averaged over onsets in either left or right V1, which resulted in an average locomotion onset response curve for left and right V1, for each mouse and condition (closed and open loop). These curves were subsequently used for calculating the correlation coefficient in the same window as above. The correlation coefficients from left and right V1 are then averaged and reported as individual data points in the figure panel. For this analysis, we excluded mice in which the responses were smaller than 1% ΔF/F in either of the two conditions (3 of 54 mice), as the correlation analysis yields noise in the absence of a clear response.

For the correlation analyses shown in **Figures 5-7, S6-S8**, we first calculated the correlation coefficient of neuronal activity (or fluorescence change, in the case of Tlx3-Cre mice that had been retro-orbitally injected with an AAV-PHP.eB-Ef1α-DIO-eGFP to express eGFP in L5 IT neurons) for all possible pairs of the 12 regions of interest. We then determined the distance between each pair of ROIs and normalized this value by the distance between bregma and lambda obtained from the images during initial surgery for each mouse (**Figure S6A**). We plotted the correlation against the distance for each pair of regions of interest and binned the data in a 40 x 40 grid and smoothed the resulting image with a gaussian filter (**Figure S6B**), to obtain the heatmaps shown in **Figures 6A and 6B, 6D and 6E, 7A and 7B, 7D and 7E, 7G and 7H, S6B-S6D, S6G and S6H, S7A and S7B, S8C and S8D**.

Contour lines in the heatmaps were then drawn to 50% of the maximum value in the plot. The boxplot quantifications (**Figures 6C, 6F, 7C, 7F, 7I, S6E, S6H, S7C, S8E**) were calculated as the change in correlation coefficient of activity after and before treatment for all pairs of regions of interest, normalized to the value before treatment and split into short- and long-range using a threshold of 0.9 of the bregma-lambda distance for each mouse. For pairwise neuronal correlations that had been recorded with two-photon imaging (**Figure S7D**), we calculated the correlation coefficient separately for all pairs of neurons within each field of view. We noticed extreme synchrony of activity of L5 IT soma recorded in 1 field of view with two-photon imaging after clozapine injection and show both distributions for all data as well as the distribution when this 1 field of view is excluded.

For **Figure S2,** onset aligned data were first averaged for each mouse and then summed in the given combinations. The SEM was calculated over mice.

To obtain the differences in the correlation between activity and locomotion after and before clozapine injection (**Figure S5D**), we first averaged all correlation coefficients in the individual sessions of a condition (closed loop, open loop, or dark) to generate one correlation coefficient per mouse and ROI, for each condition. We then calculated and plotted the difference between these correlation values after and before clozapine injection.

We used a hierarchical bootstrap (**Figures 2C and 2D**), or a bootstrap (where data were not nested, **Figure 1L and S4J**), distribution to estimate p-values. For hierarchical bootstrap, we first resampled the data (with replacement) at the level of mice, and then, from the selected mice, resampled for recording sessions. In cases where data were not nested, bootstrap samples were drawn from the distribution of data points. We then computed the mean of this bootstrap sample and repeated this 10 000 times to generate a bootstrap distribution of the mean estimate. The p-value was then defined as the proportion of bootstrap samples higher (or lower, depending on the hypothesis) than zero. For **Figures 1L and S4J**, to account for multiple comparisons, we used family-wise error correction. The significance indicators in the figure panels therefore reflect an adjusted significance threshold, αadj, according to the formula αadj = α/m, wherein α is 0.05, 0.01 or 0.001, and m denotes the number of groups (genotypes, m = 9 in both figure panels).

For widefield imaging time course data (**Figures 1F-1L, 3B and 3C, 3E and 3F, 4, 8, S1, S3A and S3C, S4, S5A, S5E**), we used a hierarchical bootstrap estimate of the mean (Saravanan et al., 2020) and the 90% confidence interval. This was done by generating 1000 bootstrap samples chosen hierarchically (first over mice, then over onsets) for each time bin (10 ms) in the plot.

To compare the similarity of closed and open loop locomotion onset responses, we used a one-way analysis of variance (ANOVA) with genotype (**Figure 1L**) or mice (**Figure S4J**) as factors (**Table S1**), respectively, followed by a bootstrap-distribution based estimate of p-values as described above.

## ACKNOWLEDGEMENTS

We thank Tingjia Lu for production of the AAV vectors, Rebecca Jordan for performing the clear skull implants used in the hemodynamic response measurements, and members of the Keller lab for discussion and support. This project has received funding from the Swiss National Science Foundation, the Novartis Research Foundation, and the European Research Council (ERC) under the European Union’s Horizon 2020 research and innovation programme (grant agreement No 865617).

## AUTHOR CONTRIBUTIONS

MH performed the experiments and analyzed the data. All authors wrote the manuscript.

## SUPPLEMENTAL FIGURES

**Figure S1.**
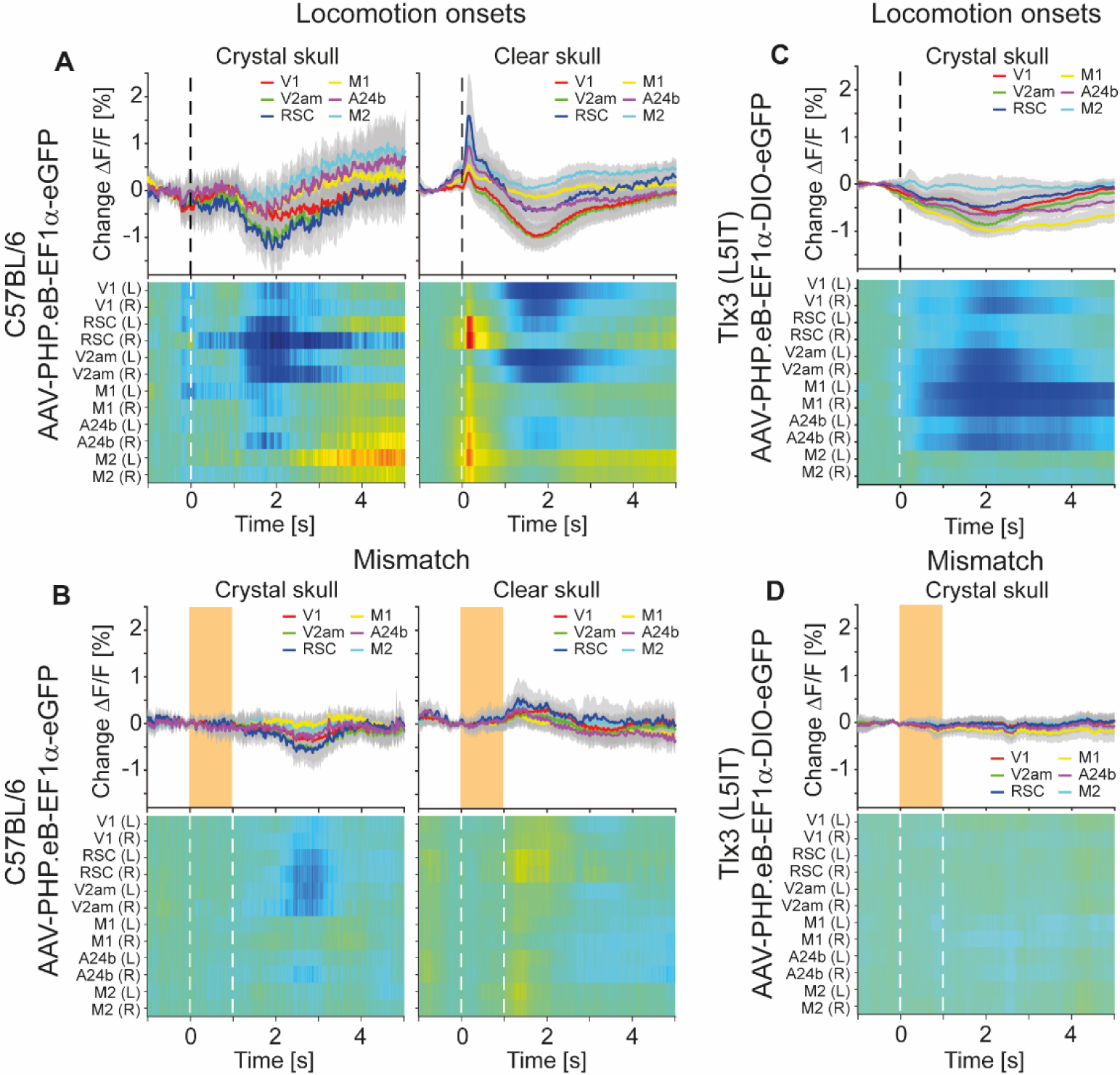
Fluorescence changes driven by hemodynamic occlusion. Related to Figure 1. (A) Average responses during locomotion onsets in C57BL/6 mice that expressed eGFP brain wide using a crystal skull preparation or in similarly transfected mice using a clear skull preparation (see Methods). Top row shows the average activity of corresponding regions in dorsal cortex. Mean (lines) and 90% confidence interval (shading) are calculated as a hierarchical bootstrap estimate for each time bin (see Methods and **Table S1**). The heatmaps in the bottom row show the responses for individual regions of interest. Heatmaps are scaled to the y-axis limits of the plot above (blue low, red high). Note, the fast onset transient apparent in the clear skull preparation was absent in the crystal skull preparation, while the slow decrease of activity was present in both. The increase in fluorescence at locomotion onset, which was primarily apparent in the clear skull preparation, is driven by hemodynamic occlusion. (B) As in **A**, but for mismatches. Orange shading (top) or white dashed lines (bottom) indicate the duration of the mismatch stimulus. (C) As in **A**, left, but for Tlx3-Cre that had been retro-orbitally injected with an AAV-PHP.eB-DIO-eGFP to express eGFP in L5 IT neurons. (D) As in **B**, left, but for Tlx3-Cre that had been retro-orbitally injected with an AAV-PHP.eB-DIO-eGFP to express eGFP in L5 IT neurons.

**Figure S2.**
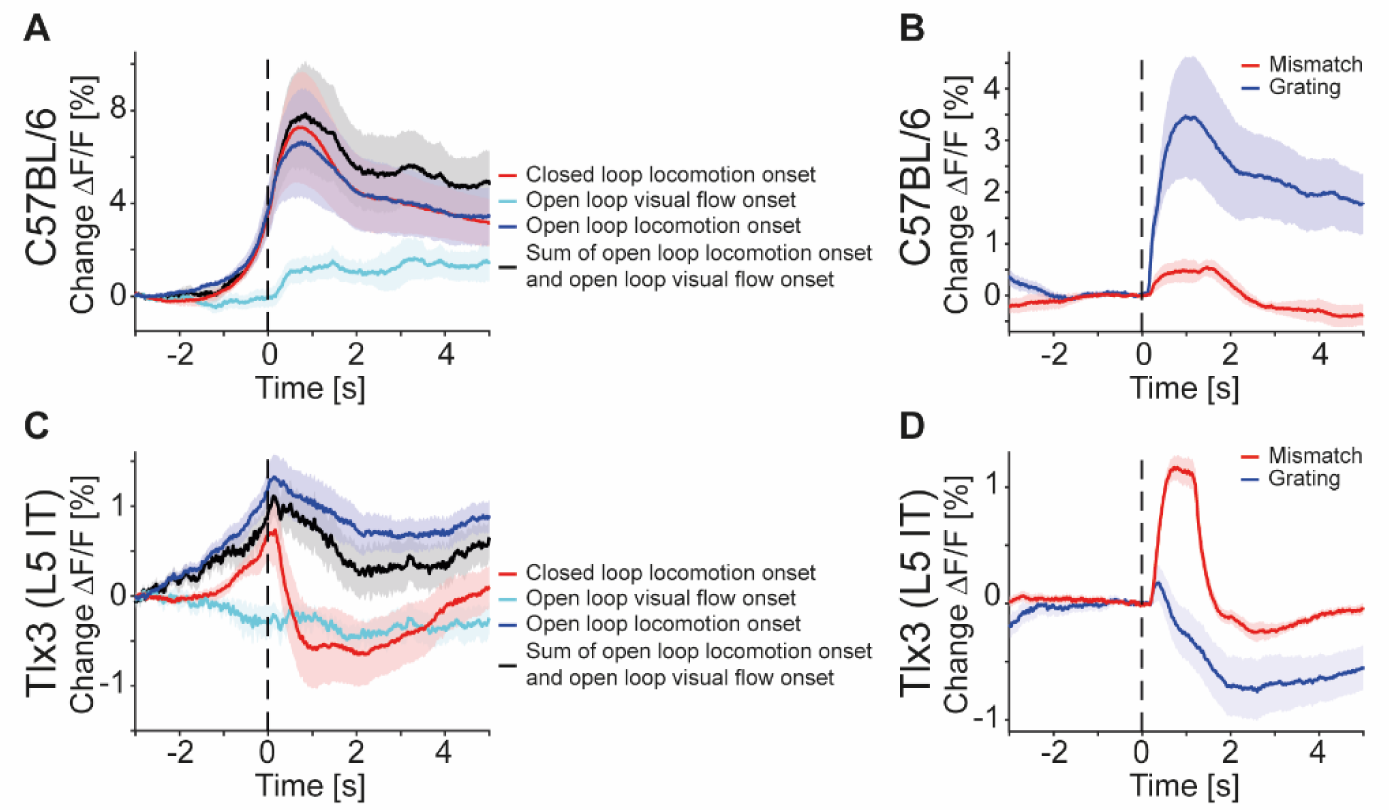
In V1, the sum of locomotion and visual flow onset could not explain the closed loop locomotion onset response of L5 IT neurons. Related to Figure 1. (A) Average responses during closed (red) and open loop (blue) locomotion onsets, as well as open loop visual flow (turquoise) onsets, and the sum of open loop locomotion and open loop visual flow onsets (black) in C57BL/6 mice that expressed GCaMP brain wide (6 mice). Closed loop locomotion onset responses in V1 were larger than open loop locomotion onset responses, and part of this difference could be explained by the visual flow onset responses. Shading indicates SEM over mice. (B) Average response during mismatch (red) and full-field drifting grating onsets (blue) in 6 C57BL/6 mice that expressed GCaMP brain wide. Same data as in Figures 3B **and 3C**. Shading indicates SEM over mice. (C) As in **A**, but for Tlx3-Cre x Ai148 mice that expressed GCaMP6 in L5 IT neurons (15 mice). Closed loop locomotion onset responses in V1 could not be explained as the sum of open loop locomotion and visual flow onset responses. (D) As in **B**, but for Tlx3-Cre x Ai148 mice that expressed GCaMP6 in L5 IT neurons (15 mice). Same data as in Figures 3E **and 3F**.

**Figure S3.**
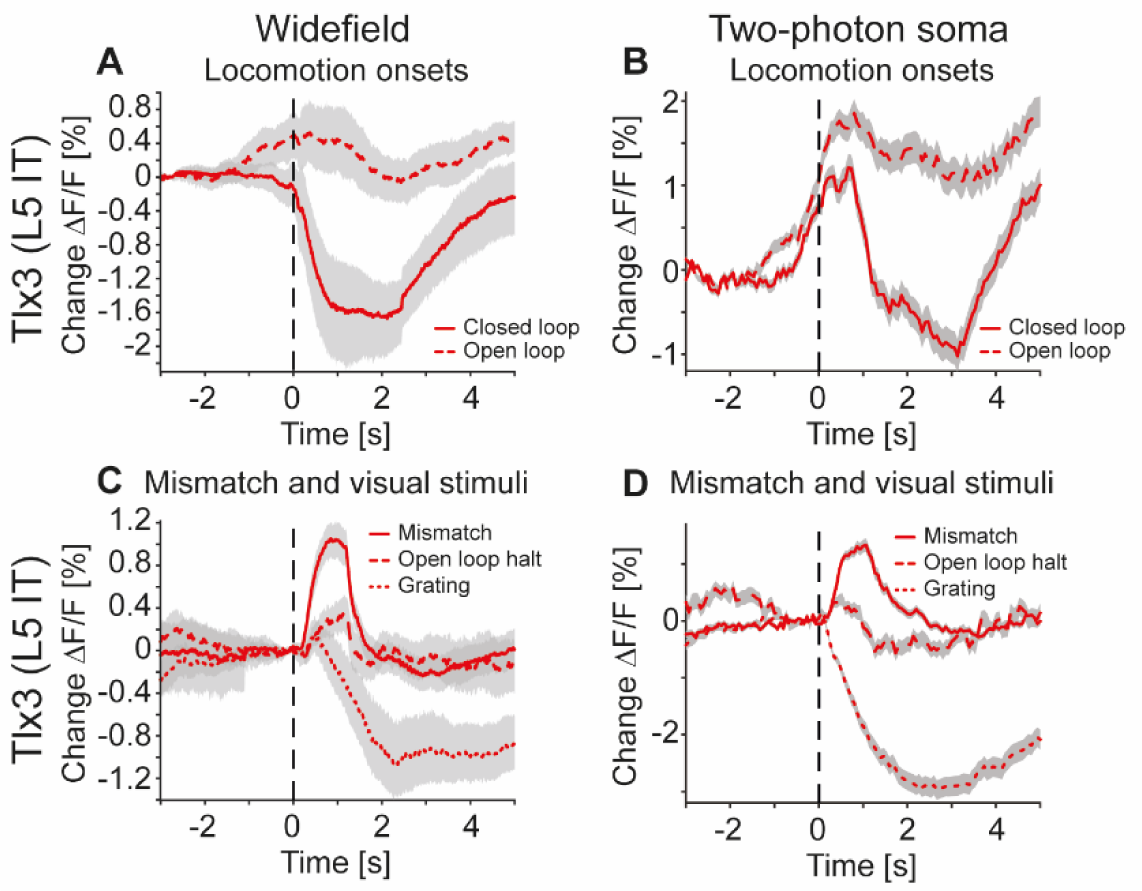
L5 IT soma recorded in V1 with two-photon imaging show similar patterns of activity as the widefield signal recorded at the surface of the dorsal cortex. Related to Figure 1. (A) Average response of V1 recorded with the widefield macroscope during closed (solid) or open loop (dashed) locomotion onsets in Tlx3-Cre x Ai148 mice that expressed GCaMP6 in L5 IT neurons. Mean (lines) and 90% confidence interval (shading) are calculated as a hierarchical bootstrap estimate for each time bin (see Methods and **Table S1**). (B) Average response of L5 soma in V1, recorded with two-photon imaging in the same Tlx3-Cre x Ai148 mice that expressed GCaMP6 in L5 IT neurons as in **A**, during either closed loop (solid) or open loop (dashed) locomotion onsets. Shading indicates SEM over 8434 neurons. (C) As in **A**, but for responses to mismatches (solid), open loop halts (dashed) or drifting grating onsets (dotted). Mean (lines) and 90% confidence interval (shading) are calculated as a hierarchical bootstrap estimate for each time bin (see Methods and **Table S1**). (D) As in **B**, but for responses to mismatches (solid), open loop halts (dashed) or drifting grating onsets (dotted). Shading indicates SEM over 8434 neurons.

**Figure S4.**
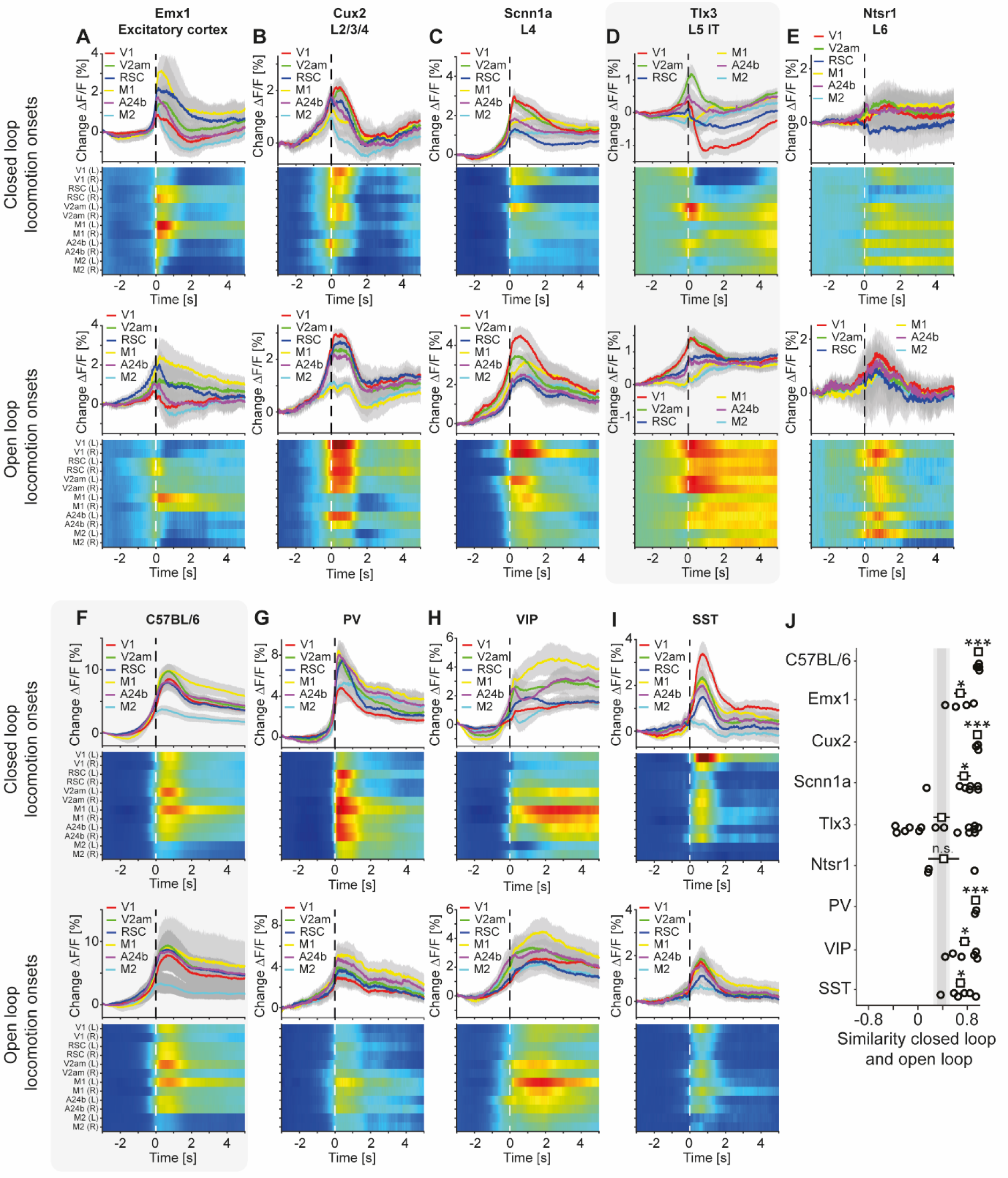
L5 IT neurons had strikingly different responses during closed and open loop locomotion onsets compared to other cortical neuron types. Related to Figure 1. (A) Average response during closed loop locomotion onsets (top) and open loop locomotion onsets (bottom) in Emx1-Cre mice that expressed GCaMP6 in excitatory cortical neurons. Mean (lines) and 90% confidence interval (shading) are calculated as a hierarchical bootstrap estimate for each time bin (see Methods and **Table S1**). Heatmaps are scaled to the y-axis limits of the plot above (blue low, red high). (B) As in **A**, but for Cux2-CreERT2 x Ai148 mice that expressed GCaMP6 in upper layer excitatory neurons. (C) As in **A**, but for Scnn1a-Cre x Ai148 mice that expressed GCaMP6 in L4 excitatory. (D) As in **A**, but for Tlx3-Cre x Ai148 mice that expressed GCaMP6 specifically in L5 IT neurons. Data are the same as in Figures 1I **and 1K**. (E) As in **A**, but for Ntsr1-Cre x Ai148 mice that expressed GCaMP6 in excitatory L6 neurons. (F) As in **A**, but for C57BL/6 mice that expressed GCaMP brain wide. Data are the same as in Figures 1F **and 1G**. (G) As in **A**, but for PV-Cre x Ai148 mice that expressed GCaMP6 in PV neurons. (H) As in **A**, but for VIP-Cre x Ai148 mice that expressed GCaMP6 in VIP neurons. (I) As in **A**, but for SST-Cre mice that expressed GCaMP6 in SST neurons. (J) Similarity of the average closed and open loop locomotion onset responses in V1 quantified as the correlation coefficient between the two in a window -5 s to +3 s around locomotion onset (left and right V1 were averaged, see Methods). Error bars indicate SEM over mice that had an average locomotion onset response of at least 1% ΔF/F in either the closed or the open loop condition (open circles: individual data points; 6 C57BL/6 mice, 4 Emx1-Cre mice, 7 Scnn1a-Cre mice, 14 Tlx3-Cre mice, 3 Ntsr1-Cre mice, 2 PV-Cre mice, 6 VIP-Cre mice, 6 Sst-Cre mice, 3 Cux2-CreERT2 mice). Statistical comparisons are against the Tlx3 data and were corrected for family-wise error rate (see Methods and **Table S1**): adjusted significance thresholds, n.s.: not significant, *: p < 0.05/9, **: p < 0.01/9, ***: p < 0.001/9. See **Table S1** for all information on statistical testing.

**Figure S5.**
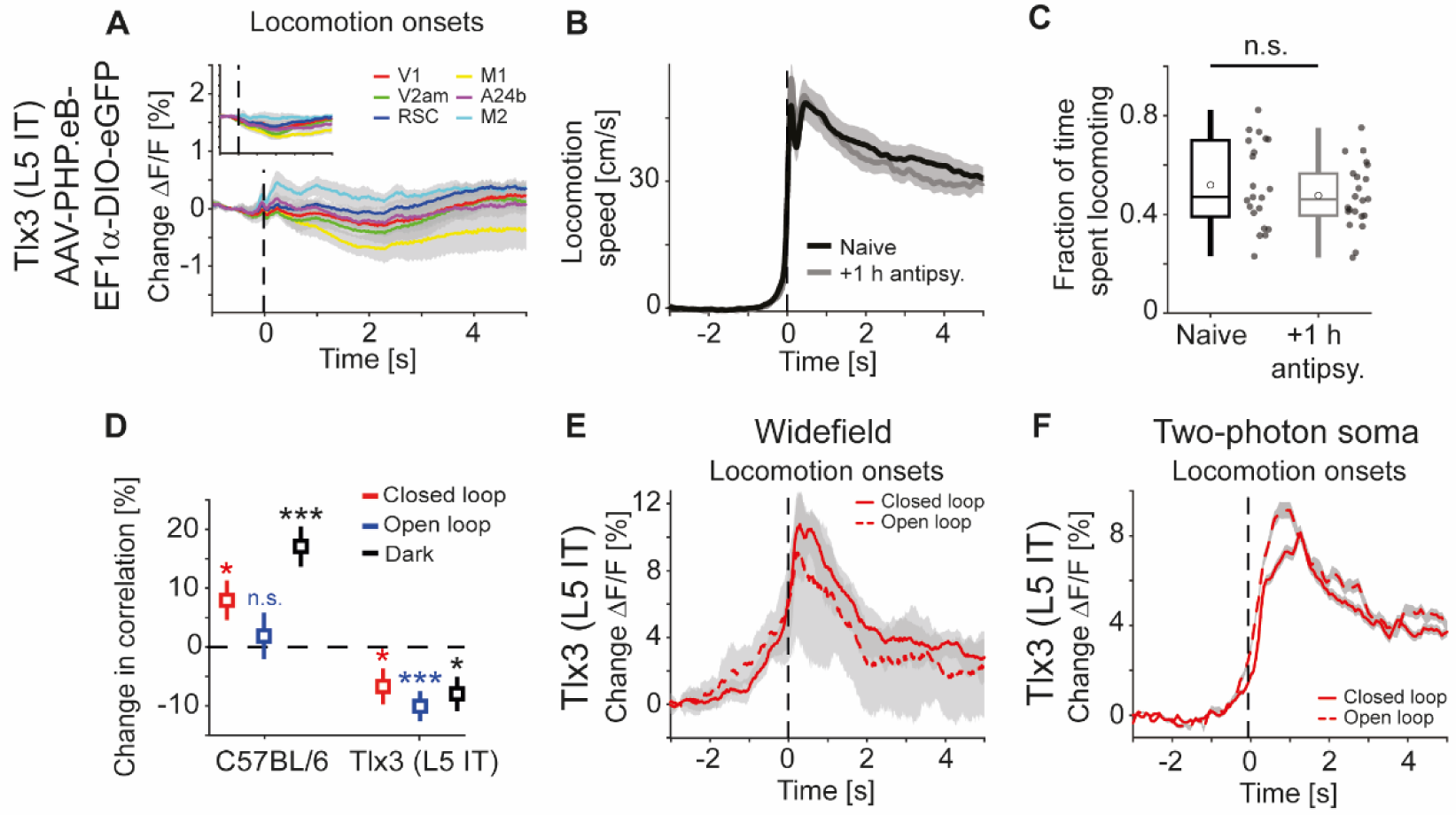
Additional information for antipsychotic drug data. Related to Figure 4. (A) Average hemodynamic response during locomotion onsets in Tlx3-Cre mice that had been retro- orbitally injected with an AAV-PHP.eB-DIO-eGFP to express eGFP in L5 IT neurons (responses were averaged across corresponding regions in both hemispheres) after a single injection of the antipsychotic drug clozapine (see Methods). Mean (lines) and 90% confidence interval (shading) are calculated as a hierarchical bootstrap estimate for each time bin (see Methods and **Table S1**). Inset: Comparison of responses to locomotion onsets in the same mice before clozapine injection (same data as in **Figure S1C**, top panel). (B) Average locomotion speed in 21 naive mice (black) and 22 mice that had been injected with either of the antipsychotic drugs clozapine, aripiprazole, or haloperidol (gray), for the data acquisition timepoint +1 h after injection. Shading indicates SEM over mice. (C) Fraction of time spent locomoting above threshold (see Methods) for the 21 naive mice (black) and 22 mice that had been injected with either of the antipsychotic drugs clozapine, aripiprazole or haloperidol, for the data acquisition timepoint +1 h after injection (gray). Boxes represent the median and the interquartile range. The open circles indicate the mean of the distributions. The whiskers mark 1.5 times the interquartile range. Dots are the individual data points. (D) Average correlation of activity and locomotion speed in dorsal cortex was increased after a single injection of the antipsychotic drug clozapine in 4 C57BL/6 mice that expressed GCaMP brain wide but decreased in 5 Tlx3-Cre x Ai148 mice that expressed GCaMP6 in L5 IT neurons, for all types of visual feedback. Error bars indicate SEM over mice and corresponding dorsal cortex regions (see Methods). n.s.: not significant, *: p < 0.05, ***: p < 0.001. See **Table S1** for all information on statistical testing. (E) Average response of V1 recorded with the widefield microscope during locomotion onsets in closed loop (solid) or open loop (dashed) in Tlx3-Cre x Ai148 mice that expressed GCaMP6 in L5 IT neurons and that had received an injection of the antipsychotic drug clozapine. Mean (lines) and 90% confidence interval (shading) are calculated as a hierarchical bootstrap estimate for each time bin (see Methods and **Table S1**). (F) As in **E**, for the same 3 Tlx3 x Ai148 mice, but for responses recorded at L5 IT soma with two- photon imaging. Shading indicates SEM over 5595 neurons.

**Figure S6.**
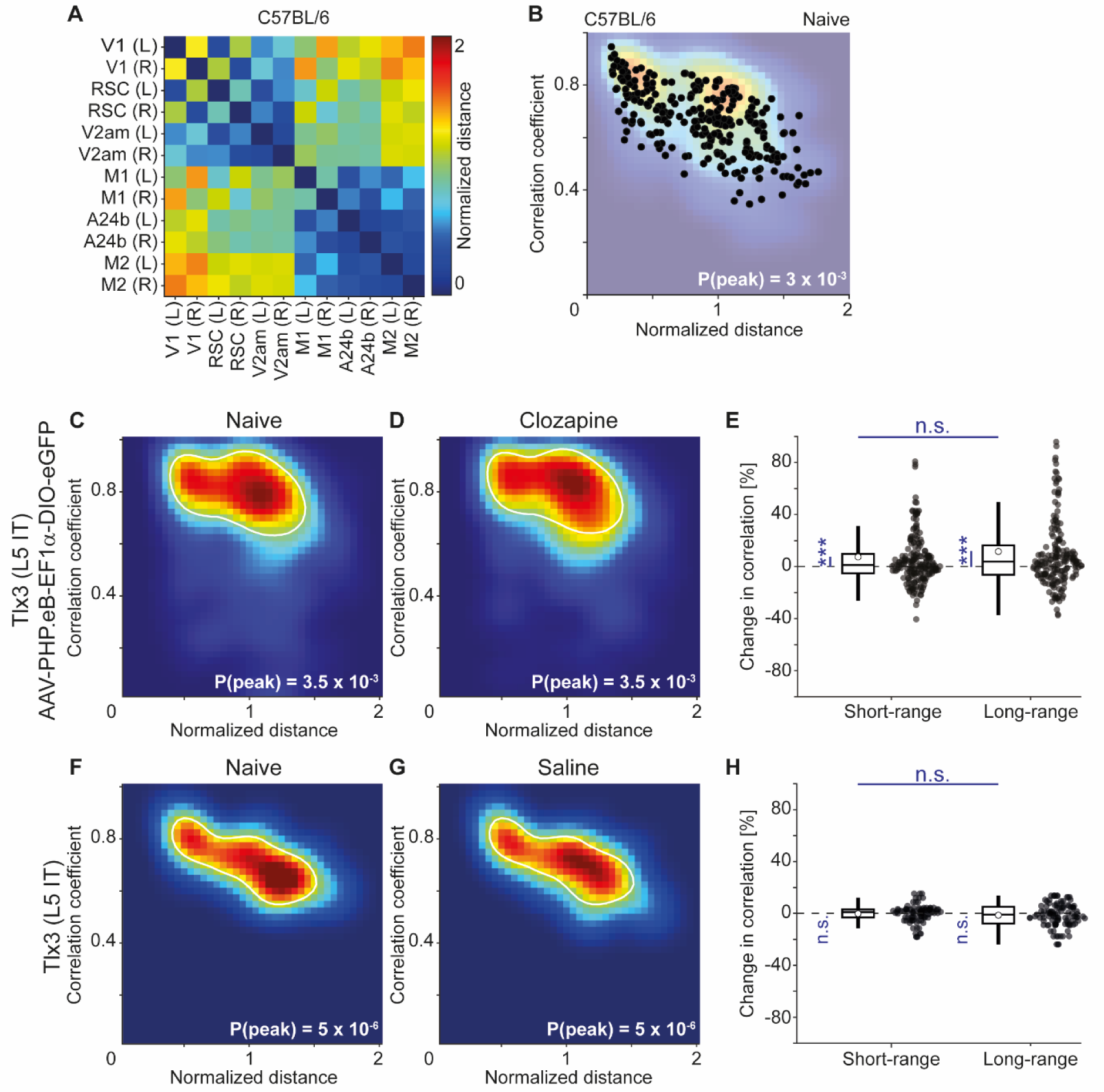
Calculation of the distance-correlation heatmaps. Related to Figure 5. (A) For each pair of dorsal cortex regions, we calculated the distance between the regions in a top- down view of dorsal cortex, normalized by the distance between bregma and lambda for each mouse (data are from an example C57BL/6 mouse that expressed GCaMP6 brain wide). (B) For each pair, we then plotted the activity correlation against this distance as calculated in **A** (black dots). We then interpolated this distribution using a 40-by-40 2D grid to obtain the density plots (faded background), shown for all mice and pairs of regions in Figures 6**, 7, and S6-S8**. (C) Density map of correlation coefficients as a function of distance between the regions, normalized across mice by the bregma-lambda distance (see panel **B** and Methods), for 6 Tlx3-Cre mice that had been retro-orbitally injected with an AAV-PHP.eB-DIO-eGFP to express eGFP in L5 IT neurons. Peak density is indicated at the bottom right of the plot and corresponds to the maximum value of the color scale (dark red). The white line is a contour line drawn at 50% of peak value. (D) As in **C**, but for the same 6 Tlx3-Cre mice that expressed eGFP in L5 IT neurons after a single injection of the antipsychotic drug clozapine. The white line is the contour from panel **C** for comparison. (E) Clozapine induced change in the fluorescence correlation between regions, normalized to the correlation coefficient in the naive state, in 6 Tlx3-Cre mice that expressed eGFP in L5 IT neurons. Data were split into short- and long-range activity correlations (see Methods). Boxes represent the median and the interquartile range. The open circle indicates the mean of the distribution. The whiskers mark 1.5 times the interquartile range. Dots are the individual data points (short-range: 204 pairs of regions, long-range: 192 pairs of regions). ***: p < 0.001, n.s.: not significant. See **Table S1** for all information on statistical testing. (F) As in **C**, but for 3 Tlx3-Cre x Ai148 mice that expressed GCaMP6 in L5 IT neurons. (G) As in **F**, but after a single injection of saline. The white line is the contour line from panel **F** for comparison. (H) Saline induced change in the activity correlation between regions, normalized to the correlation coefficient in the naive state, in 3 Tlx3-Cre x Ai148 mice. Data were split into short- and long-range activity correlations (see Methods). Boxes represent the median and the interquartile range. The open circle indicates the mean of the distribution. The whiskers mark 1.5 times the interquartile range. Dots are the individual data points (short-range: 93 pairs of regions, long-range: 105 pairs of regions). n.s.: not significant. See **Table S1** for all information on statistical testing.

**Figure S7.**
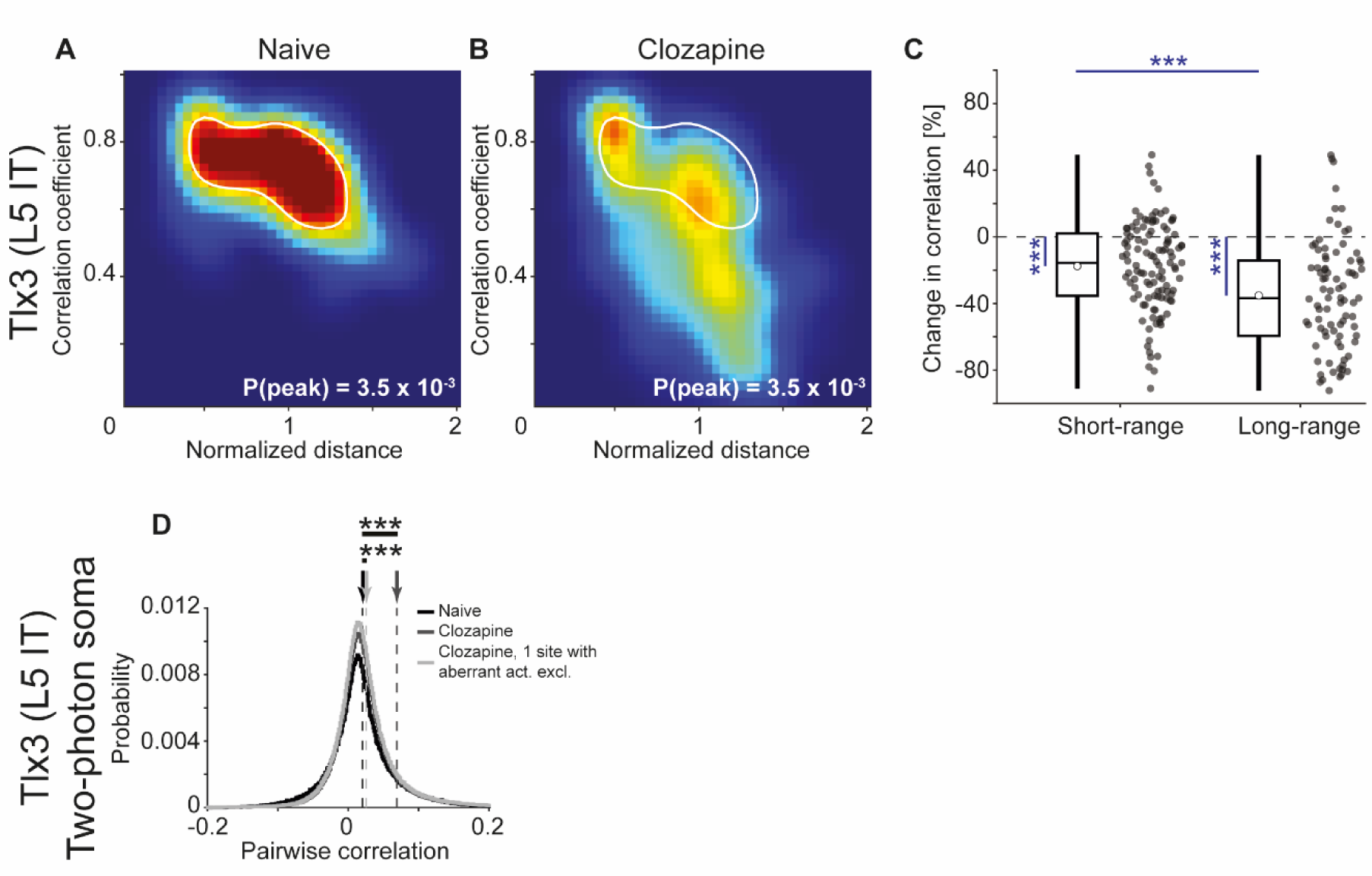
The clozapine induced decorrelation was not present in local somatic correlations. Related to Figure 6. (A) Density map of correlation coefficients as a function of distance between the regions, normalized across mice by the bregma-lambda distance (see **Figures S6A and S6B**, and Methods), for 3 Tlx3-Cre x Ai148 mice that expressed GCaMP6 in L5 IT neurons. Peak density is indicated at the bottom right of the plot and corresponds to the maximum value of the color scale (dark red). The white line is a contour line drawn at 50% of peak value. (B) As in **A**, but for the same 3 Tlx3-Cre x Ai148 mice after a single injection of the antipsychotic drug clozapine. The white line is the contour line from panel **A** for comparison. (C) Clozapine induced change in the activity correlation between regions, normalized to the correlation coefficient in the naive state, in 3 Tlx3-Cre x Ai148 mice. Data were split into short- and long-range activity correlations (see Methods). Boxes represent the median and the interquartile range. The open circle indicates the mean of the distribution. The whiskers mark 1.5 times the interquartile range. Dots are the individual data points (short-range: 114 pairs of regions, long-range: 84 pairs of regions). ***, p < 0.001. See **Table S1** for all information on statistical testing. (D) Distribution of pairwise neuronal activity correlations recorded with two-photon imaging at the L5 soma, in the same 3 Tlx3-Cre x Ai148 mice as in panels **A-C**, before (naive, black, 1991 neurons and 384 108 pairs of neurons) and after injection of the antipsychotic drug clozapine (dark gray, 7863 neurons and 885 596 pairs of neurons or pale gray, 1 field of view excluded, 7534 neurons and 831 640 pairs of neurons remaining). Dashed lines mark the mean of the respective distributions as indicated by the arrows on top. Note that after clozapine injection, we observed aberrantly high neuronal synchrony in 1 out of 25 recorded fields of view that substantially increased the average pairwise correlation as indicated by the two curves that compare the clozapine data. In either comparison, clozapine significantly increased local pairwise activity correlations.

**Figure S8.**
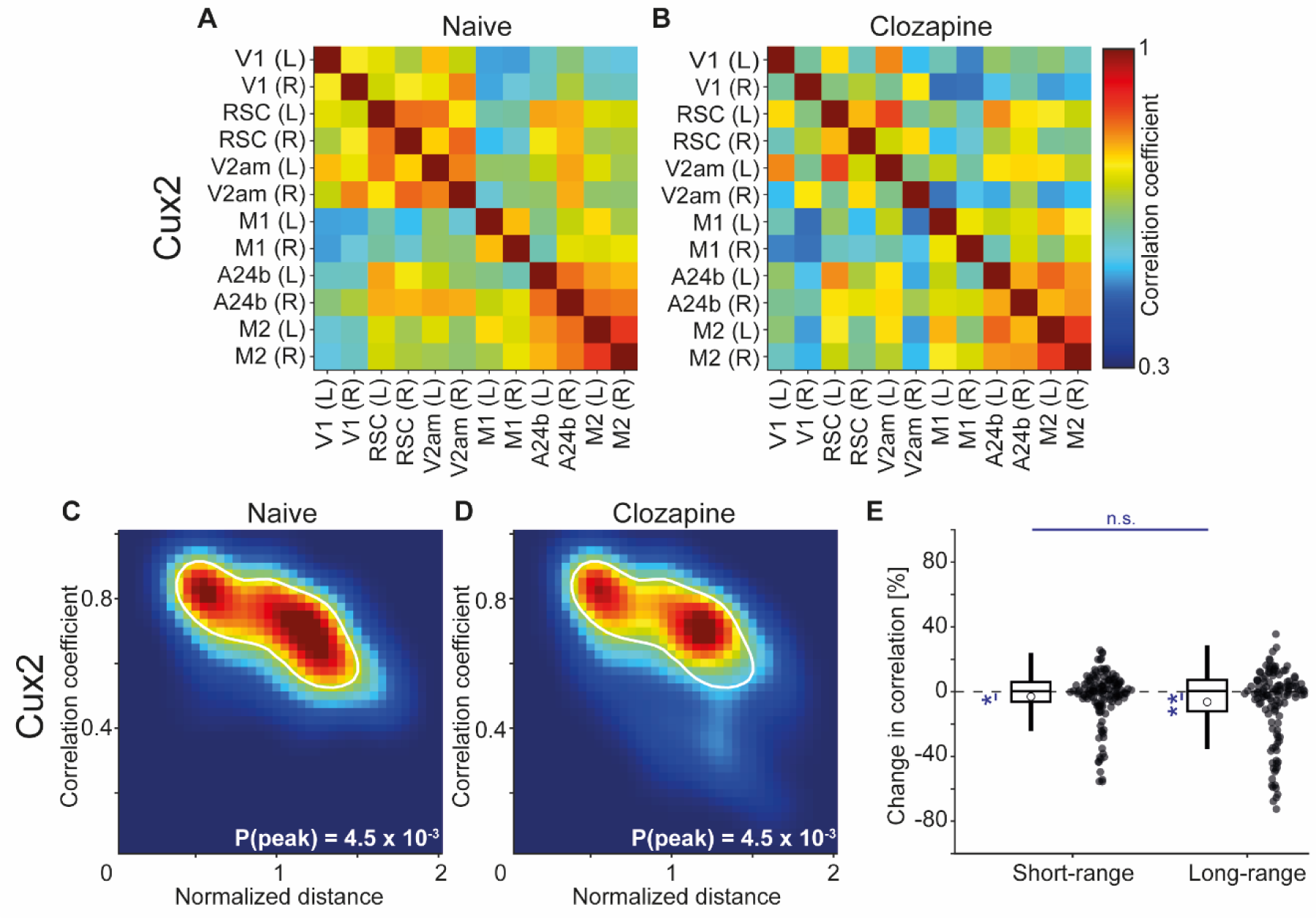
The clozapine induced decorrelation of dorsal cortex activity was weaker in superficial cortical layers than in L5. Related to Figure 6. (A) Average correlation of activity between the 12 regions of interest in 4 Cux2-CreERT2 x Ai148 mice that expressed GCaMP6 in upper layer excitatory neurons. (B) As in **A**, but for the same 4 Cux2-CreERT2 x Ai148 mice after a single injection of the antipsychotic drug clozapine. (C) Density map of correlation coefficients as a function of distance between the regions, normalized across mice by the bregma-lambda distance (see **Figures S6A and S6B**, and Methods), for 4 naive Cux2-CreERT2 x Ai148 mice that expressed GCaMP6 in upper layer excitatory neurons. Peak density is indicated at the bottom right of the plot and corresponds to the maximum value of the color scale (dark red). The white line is a contour line drawn at 50% of peak value. (D) As in **C**, but for the same 4 Cux2-CreERT2 x Ai148 mice after a single injection of the antipsychotic drug clozapine. The white line is the contour line from panel **C** for comparison. (E) Clozapine induced change in the activity correlation between regions, normalized to the correlation coefficient in the naive state, in 4 Cux2-CreERT2 x Ai148 mice. Data were split into short- and long-range activity correlations (see Methods). Boxes represent the median and the interquartile range. The open circle indicates the mean of the distribution. The whiskers mark 1.5 times the interquartile range. Dots are the individual data points (short-range: 137 pairs of regions, long-range: 127 pairs of regions). n.s.: not significant, *: p < 0.05, **: p < 0.01. See **Table S1** for all information on statistical testing.

## SUPPLEMENTAL TABLES

**Table S1.** Statistical comparisons for bar and violin plots. Note, for **Figures 1L and S4J**, the significance thresholds were corrected for multiple comparisons (see Methods).

| Figure reference | Description<br>(N<sub>1</sub> vs N<sub>2</sub>) | Test | N<sub>1</sub> (mice) | N<sub>2</sub> (mice) | Unit | P-value |
| --- | --- | --- | --- | --- | --- | --- |
| **Figure 1F** | Number of closed loop locomotion onsets in C57BL/6 mice | - | 907 (6) | - | Onsets | - |
| **Figure 1G** | Number of open loop locomotion onsets in C57BL/6 mice | - | 598 (6) | - | Onsets | - |
| **Figure 1H** | Number of open loop visual flow onsets in C57BL/6 mice | - | 416 (6) | - | Onsets | - |
| **Figure 1I** | Number of closed loop locomotion onsets in Tlx3-Cre x Ai148 mice | - | 1919 (15) | - | Onsets | - |
| **Figure 1J** | Number of open loop locomotion onsets in Tlx3-Cre x Ai148 mice | - | 1125 (15) | - | Onsets | - |
| **Figure 1K** | Number of open loop visual flow onsets in Tlx3-Cre x Ai148 mice | - | 1189 (15) | - | Onsets | - |
| **Figure 1L** | Tlx3 vs C57BL/6 | Bootstrap | 12 (15) | 12 (6) | Regions | <10<sup>-4</sup> |
|  | Tlx3 vs Emx1 | Bootstrap | 12 (15) | 12 (4) | Regions | 0.00135 |
|  | Tlx3 vs Cux2 | Bootstrap | 12 (15) | 12 (4) | Regions | <10<sup>-4</sup> |
|  | Tlx3 vs Scnn1a | Bootstrap | 12 (15) | 12 (7) | Regions | <10<sup>-4</sup> |
|  | Tlx3 vs Ntsr1 | Bootstrap | 12 (15) | 12 (3) | Regions | 0.4541 |
|  | Tlx3 vs PV | Bootstrap | 12 (15) | 12 (2) | Regions | <10<sup>-4</sup> |
|  | Tlx3 vs VIP | Bootstrap | 12 (15) | 12 (6) | Regions | 0.0001 |
|  | Tlx3 vs SST | Bootstrap | 12 (15) | 12 (5) | Regions | 0.0131 |
|  | ANOVA | ANOVA | 108 (54) | N/A | Regions | < 10<sup>-5</sup> |
| **Figure 2B** | V1 |  |  |  |  |  |
|  | Closed loop vs open loop | Hierachical bootsstrap | 308 (6) | 316 (6) | Recording session | 0.9856 |
|  | Closed loop vs dark | Hierachical bootsstrap | 308 (6) | 68 (6) | Recording session | 0.9631 |
|  | Open loop vs dark | Hierachical bootsstrap | 316 (6) | 68 (6) | Recording session | 0.8968 |
|  | RSC |  |  |  |  |  |
|  | Closed loop vs open loop | Hierachical bootsstrap | 308 (6) | 316 (6) | Recording session | 0.3423 |
|  | Closed loop vs dark | Hierachical bootsstrap | 308 (6) | 68 (6) | Recording session | 0.1492 |
|  | Open loop vs dark | Hierachical bootsstrap | 316 (6) | 68 (6) | Recording session | 0.2362 |
|  | V2am |  |  |  |  |  |
|  | Closed loop vs open loop | Hierachical bootsstrap | 308 (6) | 316 (6) | Recording session | 0.4576 |
|  | Closed loop vs dark | Hierachical bootsstrap | 308 (6) | 68 (6) | Recording session | 0.4694 |
|  | Open loop vs dark | Hierachical bootsstrap | 316 (6) | 68 (6) | Recording session | 0.4961 |
|  | M1 |  |  |  |  |  |
|  | Closed loop vs open loop | Hierachical bootsstrap | 308 (6) | 316 (6) | Recording session | 0.4417 |
|  | Closed loop vs dark | Hierachical bootsstrap | 308 (6) | 68 (6) | Recording session | 0.06 |
|  | Open loop vs dark | Hierachical bootsstrap | 316 (6) | 68 (6) | Recording session | 0.0788 |
|  | A24b |  |  |  |  |  |
|  | Closed loop vs open loop | Hierachical bootsstrap | 308 (6) | 316 (6) | Recording session | 0.8310 |
|  | Closed loop vs dark | Hierachical bootsstrap | 308 (6) | 68 (6) | Recording session | 0.2194 |
|  | Open loop vs dark | Hierachical bootsstrap | 316 (6) | 68 (6) | Recording session | 0.0846 |
|  | M2 |  |  |  |  |  |
|  | Closed loop vs open loop | Hierachical bootsstrap | 308 (6) | 316 (6) | Recording session | 0.4593 |
|  | Closed loop vs dark | Hierachical bootsstrap | 308 (6) | 68 (6) | Recording session | 0.2373 |
|  | Open loop vs dark | Hierachical bootsstrap | 316 (6) | 68 (6) | Recording session | 0.2674 |
| **Figure 2D** | V1 |  |  |  |  |  |
| | Closed loop vs open loop | Hierachical bootsstrap | 420 (15) | 394 (15) | Recording session | $<10^{-4}$ |
| | Closed loop vs dark | Hierachical bootsstrap | 420 (15) | 194 (15) | Recording session | $<10^{-4}$ |
|  | Open loop vs dark | Hierachical bootsstrap | 394 (15) | 194 (15) | Recording session | 0.2103 |
|  | RSC |  |  |  |  |  |
| | Closed loop vs open loop | Hierachical bootsstrap | 420 (15) | 394 (15) | Recording session | $<10^{-4}$ |
|  | Closed loop vs dark | Hierachical bootsstrap | 420 (15) | 194 (15) | Recording session | 0.0001 |
|  | Open loop vs dark | Hierachical bootsstrap | 394 (15) | 194 (15) | Recording session | 0.0021 |
|  | V2am |  |  |  |  |  |
| | Closed loop vs open loop | Hierachical bootsstrap | 420 (15) | 394 (15) | Recording session | $<10^{-4}$ |
| | Closed loop vs dark | Hierachical bootsstrap | 420 (15) | 194 (15) | Recording session | $<10^{-4}$ |
|  | Open loop vs dark | Hierachical bootsstrap | 394 (15) | 194 (15) | Recording session | 0.1337 |
|  | M1 |  |  |  |  |  |
| | Closed loop vs open loop | Hierachical bootsstrap | 420 (15) | 394 (15) | Recording session | $<10^{-4}$ |
|  | Closed loop vs dark | Hierachical bootsstrap | 420 (15) | 194 (15) | Recording session | 0.3115 |
|  | Open loop vs dark | Hierachical bootsstrap | 394 (15) | 194 (15) | Recording session | 0.0002 |
|  | A24b |  |  |  |  |  |
| | Closed loop vs open loop | Hierachical bootsstrap | 420 (15) | 394 (15) | Recording session | $<10^{-4}$ |
|  | Closed loop vs dark | Hierachical bootsstrap | 420 (15) | 194 (15) | Recording session | 0.3511 |
|  | Open loop vs dark | Hierachical bootsstrap | 394 (15) | 194 (15) | Recording session | 0.0001 |
|  | M2 |  |  |  |  |  |
|  | Closed loop vs open loop | Hierachical bootsstrap | 420 (15) | 394 (15) | Recording session | 0.0005 |
|  | Closed loop vs dark | Hierachical bootsstrap | 420 (15) | 194 (15) | Recording session | 0.2686 |
|  | Open loop vs dark | Hierarchical bootstrap | 394 (15) | 194 (15) | Recording session | 0.0156 |
| **Figure 3B** | Number of mismatch onsets in C57BL/6 mice | - | 2512 (6) | - | Onsets | - |
| **Figure 3C** | Number of grating onsets in C57BL/6 mice | - | 1858 (6) | - | Onsets | - |
| **Figure 3E** | Number of mismatch onsets in Tlx3-Cre x Ai148 mice | - | 3297 (15) | - | Onsets | - |
| **Figure 3F** | Number of mismatch onsets in Tlx3-Cre x Ai148 mice | - | 5318 (15) | - | Onsets | - |
| **Figure 4A** | Number of closed loop locomotion onset in clozapine treated C57BL/6 mice | - | 273 (4) | - | Onsets | - |
| **Figure 4A, inset** | Number of closed loop locomotion onset in naïve C57BL/6 mice | - | 720 (4) | - | Onsets | - |
| **Figure 4B** | Number of open loop locomotion onset in clozapine treated C57BL/6 mice | - | 145 (4) | - | Onsets | - |
| **Figure 4B, inset** | Number of open loop locomotion onset in naïve C57BL/6 mice | - | 570 (4) | - | Onsets | - |
| **Figure 4C** | Number of open loop visual flow onsets in clozapine treated C57BL/6 mice that were not locomoting | - | 99 (4) | - | Onsets | - |
| **Figure 4C, inset** | Number of open loop visual flow onsets in naïve C57BL/6 mice that were not locomoting | - | 404 (4) | - | Onsets | - |
| **Figure 4D** | Number of closed loop locomotion onset in clozapine treated Tlx3-Cre x Ai148 mice | - | 707 (5) | - | Onsets | - |
| **Figure 4D, inset** | Number of closed loop locomotion onset in naïve Tlx3-Cre x Ai148 mice | - | 1101 (5) | - | Onsets | - |
| **Figure 4E** | Number of open loop locomotion onset in clozapine treated Tlx3-Cre x Ai148 mice | - | 350 (5) | - | Onsets | - |
| **Figure 4E, inset** | Number of open loop locomotion onset in naïve Tlx3-Cre x Ai148 mice | - | 348 (5) | - | Onsets | - |
| **Figure 4F** | Number of open loop visual flow onsets in clozapine treated Tlx3-Cre x Ai148 mice that were not locomoting | - | 514 (5) | - | Onsets | - |
| **Figure 4F, inset** | Number of open loop visual flow onsets in naïve Tlx3-Cre x Ai148 mice that were not locomoting | - | 568 (5) | - | Onsets | - |
| **Figure 6C** | Short-range vs no change | T-test | 122 (4) | - | Pairs of regions | 0.3791 |
|  | Long-range vs no change | T-test | 142 (4) | - | Pairs of regions | 0.9035 |
|  | Short-range vs long-range | Ranksum | 122 (4) | 142 (4) | Pairs of regions | 0.0511 |
| **Figure 6F** | Short-range vs no change | T-test | 182 (5) | - | Pairs of regions | <10<sup>-5</sup> |
|  | Long-range vs no change | T-test | 148 (5) | - | Pairs of regions | <10<sup>-10</sup> |
|  | Short-range vs long-range | Ranksum | 182 (5) | 148 (5) | Pairs of regions | <10<sup>-4</sup> |
| **Figure 7C** | Short-range vs no change | T-test | 114 (3) | - | Pairs of regions | <10<sup>-5</sup> |
|  | Long-range vs no change | T-test | 84 (3) | - | Pairs of regions | <10<sup>-7</sup> |
|  | Short-range vs long-range | Ranksum | 114 (3) | 84 (3) | Pairs of regions | 0.0347 |
| **Figure 7F** | Short-range vs no change | T-test | 114 (3) | - | Pairs of regions | 0.7058 |
|  | Long-range vs no change | T-test | 84 (3) | - | Pairs of regions | 0.0435 |
|  | Short-range vs long-range | Ranksum | 114 (3) | 84 (3) | Pairs of regions | 0.0199 |
| **Figure 7I** | Short-range vs no change | T-test | 114 (3) | - | Pairs of regions | 0.4865 |
|  | Long-range vs no change | T-test | 84 (3) | - | Pairs of regions | 0.2184 |
|  | Short-range vs long-range | Ranksum | 114 (3) | 84 (3) | Pairs of regions | 0.1633 |
| **Figure 8A** | Number of mismatch onsets in naïve Tlx3-Cre x Ai148 mice | - | 2464 (11) | - | Onsets | - |
|  | Number of mismatch onsets in Tlx3-Cre x Ai148 mice treated with antipsychotic drugs | - | 2017 (11) | - | Onsets | - |
| **Figure 8B** | Number of grating onsets in naïve Tlx3-Cre x Ai148 mice | - | 3942 (11) | - | Onsets | - |
|  | Number of grating onsets in Tlx3-Cre x Ai148 mice treated with antipsychotic drugs | - | 1645 (11) | - | Onsets | - |
| **Figure S1A, left** | Number of locomotion onsets in C57BL/6 mice that expressed GFP brain wide and were implanted with crystal skulls | - | 96 (3) | - | Onsets | - |
| **Figure S1A, right** | Number of locomotion onsets in C57BL/6 mice that expressed GFP brain wide with clear skull cranial windows | - | 615 (5) | - | Onsets | - |
| **Figure S1B, left** | Number of mismatch onsets in C57BL/6 mice that expressed GFP brain wide and were implanted with crystal skulls | - | 229 (3) | - | Onsets | - |
| **Figure S1B, right** | Number of mismatch onsets in C57BL/6 mice that expressed GFP brain wide with clear skull cranial windows | - | 228 (5) | - | Onsets | - |
| **Figure S1C** | Number of locomotion onsets in Tlx3-Cre mice that expressed GFP in L5 IT neurons | - | 1880 (7) | - | Onsets | - |
| **Figure S1D** | Number of mismatch onsets in Tlx3-Cre mice that expressed GFP in L5 IT neurons | - | 451 (7) | - | Onsets | - |
| **Figure S3A** | Number of closed loop locomotion onsets in Tlx3-Cre x Ai148 mice recorded with the widefield microscope | - | 1012 (7) | - | Onsets | - |
|  | Number of open loop locomotion onsets in Tlx3-Cre x Ai148 mice recorded with the widefield microscope | - | 678 (7) | - | Onsets | - |
| **Figure S3C** | Number of mismatch onsets in Tlx3-Cre x Ai148 mice recorded with the widefield microscope | - | 1038 (7) | - | Onsets | - |
|  | Number of open loop halts in Tlx3-Cre x Ai148 mice recorded with the widefield microscope | - | 410 (7) | - | Onsets | - |
|  | Number of grating onsets in Tlx3-Cre x Ai148 mice recorded with the widefield microscope | - | 2014 (7) | - | Onsets | - |
| **Figure S4A** | Number of closed loop locomotion onsets in Emx1-Cre mice | - | 438 (4) | - | Onsets | - |
|  | Number of open loop locomotion onsets in Emx1-Cre mice | - | 286 (4) | - | Onsets | - |
| **Figure S4B** | Number of closed loop locomotion onsets in Cux2-CreERT2 x Ai148 mice | - | 415 (4) | - | Onsets | - |
|  | Number of open loop locomotion onsets in Cux2-CreERT2 x Ai148 mice | - | 433 (4) | - | Onsets | - |
| **Figure S4C** | Number of closed loop locomotion onsets in Scnn1a x Ai148 mice | - | 839 (7) | - | Onsets | - |
|  | Number of open loop locomotion onsets in Scnn1a x Ai148 mice | - | 313 (7) | - | Onsets | - |
| **Figure S4D** | Number of closed loop locomotion onsets in Tlx3-Cre x Ai148 mice | - | 1919 (15) | - | Onsets | - |
|  | Number of open loop locomotion onsets in Tlx3-Cre x Ai148 mice | - | 1125 (15) | - | Onsets | - |
| **Figure S4E** | Number of closed loop locomotion onsets in Ntsr1-Cre x Ai148 mice | - | 368 (3) | - | Onsets | - |
|  | Number of open loop locomotion onsets in Ntsr1-Cre x Ai148 mice | - | 112 (3) | - | Onsets | - |
| **Figure S4F** | Number of closed loop locomotion onsets in C57BL/6 mice | - | 907 (6) | - | Onsets | - |
|  | Number of open loop locomotion onsets in C57BL/6 mice | - | 598 (6) | - | Onsets | - |
| **Figure S4G** | Number of closed loop locomotion onsets in PV-Cre x Ai148 mice | - | 236 (2) | - | Onsets | - |
|  | Number of open loop locomotion onsets in PV-Cre x Ai148 mice | - | 110 (2) | - | Onsets | - |
| **Figure S4H** | Number of closed loop locomotion onsets in VIP-Cre x Ai148 mice | - | 618 (6) | - | Onsets | - |
|  | Number of open loop locomotion onsets in VIP-Cre x Ai148 mice | - | 376 (6) | - | Onsets | - |
| **Figure S4I** | Number of closed loop locomotion onsets in Sst-Cre x Ai148 mice | - | 747 (5) | - | Onsets | - |
|  | Number of open loop locomotion onsets in Sst-Cre x Ai148 mice | - | 558 (5) | - | Onsets | - |
| **Figure S4J** | Tlx3 vs C57BL/6 | Bootstrap | 14 | 6 | V1 / Mice | <10<sup>-4</sup> |
|  | Tlx3 vs Emx1 | Bootstrap | 14 | 4 | V1 / Mice | 0.0042 |
|  | Tlx3 vs Cux2 | Bootstrap | 14 | 3 | V1 / Mice | <10<sup>-4</sup> |
|  | Tlx3 vs Scnn1a | Bootstrap | 14 | 7 | V1 / Mice | 0.0042 |
|  | Tlx3 vs Ntsr1 | Bootstrap | 14 | 3 | V1 / Mice | 0.4087 |
|  | Tlx3 vs PV | Bootstrap | 14 | 2 | V1 / Mice | <10<sup>-4</sup> |
|  | Tlx3 vs VIP | Bootstrap | 14 | 6 | V1 / Mice | 0.0021 |
|  | Tlx3 vs SST | Bootstrap | 14 | 6 | V1 / Mice | 0.0032 |
|  | ANOVA | ANOVA | 54 | N/A | V1 / Mice | 0.0004 |
| **Figure S5A** | Number of locomotion onsets in Tlx3-Cre mice | - | 2331 (7) | - | Onsets | - |
| **Figure S5C** | Naive vs +1 h antipsy. | Ranksum | 21 | 22 | Mice | 0.4735 |
| **Figure S5D** | C57BL/6 |  |  |  |  |  |
|  | Closed loop vs no change | T-test | 24 (4) | - | Regions | 0.0277 |
|  | Open loop vs no change | T-test | 24 (4) | - | Regions | 0.6384 |
|  | Dark vs no change | T-test | 24 (4) | - | Regions | <10<sup>-4</sup> |
|  | Tlx3 |  |  |  |  |  |
|  | Closed loop vs no change | T-test | 30 (5) | - | Regions | 0.0365 |
|  | Open loop vs no change | T-test | 30 (5) | - | Regions | 0.0004 |
|  | Dark vs no change | T-test | 30 (5) | - | Regions | 0.0104 |
| **Figure S5E** | Number of closed loop locomotion onsets in Tlx3-Cre x Ai148 mice | - | 376 (3) | - | Onsets | - |
|  | Number of open loop locomotion onsets in Tlx3-Cre x Ai148 mice | - | 245 (3) | - | Onsets | - |
| **Figure S6E** | Short-range vs no change | T-test | 204 (7) | - | Pairs of regions | 0.0006 |
|  | Long-range vs no change | T-test | 192 (7) | - | Pairs of regions | <10<sup>-4</sup> |
|  | Short-range vs long-range | Ranksum | 204 (7) | 192 (7) | Pairs of regions | 0.2540 |
| **Figure S6H** | Short-range vs no change | T-test | 93 (3) | - | Pairs of regions | 0.7884 |
|  | Long-range vs no change | T-test | 105 (3) | - | Pairs of regions | 0.1176 |
|  | Short-range vs long-range | Rank-sum | 93 (3) | 105 (3) | Pairs of regions | 0.2319 |
| **Figure S7C** | Short-range vs no change | T-test | 114 (3) | - | Pairs of regions | $<10^{-4}$ |
| | Long-range vs no change | T-test | 84 (3) | - | Pairs of regions | $<10^{-4}$ |
| | Short-range vs long-range | Ranksum | 114 (3) | 84 (3) | Pairs of regions | $<10^{-4}$ |
| **Figure S7D** | Naive vs clozapine | Ranksum | 384108 (7) | 885596 | Pairs of neurons | $<10^{-4}$ |
| | Naive vs clozapine 1 site excl. | Ranksum | 384108 (7) | 831640 | Pairs of neurons | $<10^{-4}$ |
| **Figure S8E** | Short-range vs no change | T-test | 137 (4) | - | Pairs of regions | 0.0262 |
|  | Long-range vs no change | T-test | 127 (4) | - | Pairs of regions | 0.0019 |
|  | Short-range vs long-range | Ranksum | 137 (4) | 127 (4) | Pairs of regions | 0.9537 |

**Table S2.** Indicator expression strategy in individual mice.

| Nbr. mice | Genotype | Virus | Figures |
| --- | --- | --- | --- |
| 3 | C57BL/6 | AAV-PHP.eB-hSyn1-jGCaMP7f | 1C, 1F-1H, 1L, 2A and 2B, 3A-3C, 4A-4C, 5A and 5B, 6A-6C, S2, S4F, S4J, S5B-S5D, S6A and S6B |
| 3 | C57BL/6 | AAV-PHP.eB-EF1α-GCaMP6s | 1F-1H, 1L, 2B, 3B and 3C, 4A-4C, 5A and 5B, 6A-6C, S2, S4F, S4J, S5B-S5D |
| 8 | C57BL/6 | AAV-PHP.eB-EF1α-eGFP | S1 |
| 4 | Emx1-Cre | AAV-PHP.eB-DIO-EF1α-GCaMP6s | 1L, S4A and S4J |
| 4 | Cux2-CreERT2 x Ai148 | - | 1L, S4B, S4J and S8 |
| 7 | Scnn1a-Cre x Ai148 | - | 1L, S4C and S4J |
| 25 | Tlx3-Cre x Ai148 | - | 1D, 1I-1L, 2C and 2D, 3D-3F, 4D-4F, 5C and 5D, 6D-6F, 7, 8, S2, S3D, S4D, S4J, S5B-S5F, S6G-S6H, S7 |
| 7 | Tlx3-Cre | AAV-PHP.eB-EF1α-DIO-eGFP | S1C and S1D, S5A, S6C-S6E |
| 3 | Ntsr1-Cre x Ai148 | - | 1L, S4E and S4J |
| 2 | PV-Cre x Ai148 | - | 1L, S4G and S4J |
| 6 | VIP-Cre x Ai148 | - | 1L, S4H and S4J |
| 4 | SST-Cre | AAV-PHP.eB-DIO-EF1α-jGCaMP7f | 1L, S4I and S4J |
| 1 | SST-Cre x Ai148 | - | 1L, S4I and S4J |

